# HDAC6 inhibition rewires carbon metabolism to expose an asparagine dependency in multiple myeloma

**DOI:** 10.64898/2026.11.25.768827

**Authors:** Matthew Hiller, Cian Darcy, Izabella Cymer, Sandra Franco-Caspueñas, Joanna Fay, Tanya Schloms, Keelan Farnan, Yu Wang, Stephanie Gilmore, Ludovica Di Martino, Andrew Roe, Kieran Wynne, Nora Rauch, Emma Kerr, Heiko Düssmann, Massimiliano Garre, Catriona Dowling, Marie McIlroy, Jochen H.M Prehn, John Quinn, Siobhan V. Glavey, Daniel A. Rodrigues, Ann M. Hopkins, Martina Wallace, Tríona Ní Chonghaile

**Affiliations:** Department of Physiology and Medical Physics, RCSI University of Medicine and Health Sciences, Dublin, Ireland; Department of Surgery, RCSI University of Medicine and Health Sciences, Dublin, Ireland; Department of Pathology, RCSI University of Medicine and Health Sciences, Dublin, Ireland; Neurobiology of Hearing and Myelinopathies Group, Institute for Biomedical Research Sols-Morreale (IIBM), Spanish National Research Council (CSIC), Autonomous University of Madrid (UAM), Madrid, Spain; Department of Medicine, RCSI University of Medicine and Health Sciences, Dublin, Ireland; Systems Biology Ireland, University College Dublin, Dublin, Ireland; Johnston Cancer Research Centre, Queen’s University, Belfast, N. Ireland; Super Resolution Imaging Consortium (SRIC), Royal College of Surgeons in Ireland, Dublin; Department of Haematology and Pathology, Beaumont Hospital, Dublin, Ireland; Department of Chemistry, RCSI University of Medicine and Health Sciences, Dublin, Ireland; Conway Institute, University College Dublin, Dublin, Ireland

**Keywords:** HDAC6, Multiple Myeloma, glycolysis, Asparagine, L-ASN, CAM

## Abstract

Multiple myeloma (MM) remains a challenging malignancy with limited therapeutic options upon relapse/refractory disease, necessitating novel combination strategies. Malignant plasma cells sustain continuous immunoglobulin production, a biosynthetic and secretory burden that places heavy demand on central carbon metabolism and creates potentially targetable metabolic vulnerabilities. Previously, we found a role for HDAC6 in modulating glycolytic metabolism. Our goal here was to find induced metabolic vulnerabilities that could be targeted after HDAC6 inhibition or knockdown. Genetic silencing of HDAC6 or pharmacological inhibition with the selective inhibitor BAS-2, suppressed glycolytic flux, reduced pyruvate and lactate labelling from ^13^C_6_-glucose, and remodeled mitochondrial morphology in MM cell lines. Global proteomic analysis revealed coordinated upregulation of amino acid transporters (SLC3A2, SLC7A1, SLC1A5, SLC7A5), and spent-medium measurements confirmed an increased depletion of extracellular amino acids. Collectively, this data indicates that BAS-2-treated cells increased their dependence on amino acid import. To identify essential amino acids, we screened MM cells treated with BAS-2 and individually depleted each of the 20 proteinogenic amino acids. Upon depletion of either asparagine, arginine, or aspartate enhanced BAS-2 induced cell death. Combination treatment of BAS-2 and the FDA-approved drug L-asparaginase (L-ASNase), which depletes asparagine, demonstrated synergistic cytotoxicity across a panel of MM cell lines. The combination treatment induces a reliance on asparagine synthesis from pyruvate, as identified by labelling from ^13^C_6_-glucose. Mechanistically the combination induced inhibition of protein synthesis, activated the integrated stress response, and increased ATF4 and CHOP protein expression. Lastly, we showed *in vivo* in the chick chorioallantoic membrane (CAM) model that BAS-2 and L-ASNase reduced tumour volume and CD138 positivity. In summary, these findings establish HDAC6 inhibition as a translatable metabolic target which rewires amino acid import that can be exploited with L-ASNase in MM.

## Main

Multiple myeloma (MM) is a malignancy of mature plasma B cells that develops in the bone marrow. Despite the development of novel therapeutic approaches, including proteasome inhibitors, immunomodulatory agents, antibody-drug conjugates and CAR-T cell therapy, MM inevitably relapses. Relapsed/refractory MM becomes more difficult to treat on each relapse, and currently it is still an incurable disease, highlighting the need for novel therapeutic paradigms^1–5^. A defining feature of MM biology is the continuous secretion of globular IgG-like proteins, called M-Proteins, that induces a high metabolic and biomass demand^6,7^. This high metabolic demand is met through catabolising carbon sources, such as glucose through glycolysis and the tricarboxylic acid (TCA) cycle, as well as oxidizing glutamine through glutaminolysis or fatty acids through β-oxidation^8–12^. Beyond generating energy, these metabolic pathways generate resources such as amino acids for protein production and nucleotides for DNA replication and biomass generation. Although elegant studies have defined the metabolic rewiring of MM cells, their metabolic plasticity means it remains unclear whether direct or indirect perturbation of metabolic pathways can be exploited therapeutically.

Despite intense research efforts, few metabolic inhibitors have transitioned to the clinic for the treatment of cancer. Antimetabolites targeting folate metabolism or purine analogues, together with L-asparaginase, which depletes circulating asparagine, remain the principal examples and have been in clinical use for decades^13^. L-ASNase is a standard-of-care therapeutic strategy in paediatric B cell acute lymphoblastic leukaemia (B-ALL), and has been clinically tested in a variety of other haematological malignancies^14,15^. Inhibitors of mutant isocitrate dehydrogenase are a more recent exception^16,17^, and the metabolic dependencies of most tumours, particularly those in central carbon metabolism, have yet to be successfully exploited therapeutically.

Histone Deacetylases (HDAC) are a family of enzymes whose role is to remove acetyl groups from lysine residues, affecting the activity of both nuclear and cytoplasmic proteins. Inhibitors of the HDAC family are clinically approved in the treatment of MM^18–20^. HDAC6 is an interesting member of the family, as it is cytosolically located and has deacetylase activity and a ubiquitin-binding domain that shuttles unfolded proteins to the aggresome^21^. Given HDAC6’s additional, non-catalytic roles, our group has developed TTCP10, a HDAC6-targeting proteolysis targeting antigenic chimera (PROTAC) for more thorough enzyme characterisation^22^. Prior work has shown that MM is sensitive to HDAC6 inhibition (HDAC6i) particularly in combination with bortezomib^23,24^. Recently, our group identified a role for the cytoplasmic deacetylase HDAC6 in the modulation of cellular metabolism^25,26^. We previously demonstrated that HDAC6 reduces glycolytic metabolism, in triple negative breast cancer (TNBC) and non-small cell lung cancer (NSCLC) models^25,26^. Pharmacologic inhibition of HDAC6 with the selective inhibitor BAS-2, also altered the activity of fumarate hydratase, causing rewiring in TCA cycle dynamics^26^. A similar effect of HDAC6i on mitochondrial structure was also found in macrophages^27^. BAS-2 is activated on HDAC6 through a mechanism-based process that requires a specific residue, N567, in the human enzyme^22^. It remains unclear, however, whether pharmacological or genetic loss of HDAC6 function alters MM metabolism.

Given HDAC6’s newly established role in metabolic regulation, our aim was to assess if HDAC6 inhibition or knockdown alters MM metabolic programs and induces alternative means of sustaining nutrient pools, thereby revealing clinically actionable vulnerabilities. Global proteomic assessment indicated an increase in amino acid transporter expression and complementary upregulation in amino acid import. An amino acid depletion screen showed that BAS-2 treated MM cells become reliant on exogenous asparagine, arginine, and aspartate among others. Pharmacological asparagine depletion using L-ASNase synergises with BAS-2 across MM and three-dimensional bioprinted bone marrow models. L-ASNase increases cellular reliance on asparagine synthesis from glucose, a pathway restricted by BAS-2 treatment. In the chorioallantoic membrane (CAM) xenograft model, co-treatment reduced tumour burden and triggered ISR activation. These data highlight the potential translation of an induced metabolic vulnerability in multiple myeloma.

## Results

### 2.1 HDAC6 inhibition and knockdown suppresses glycolytic flux in MM

To determine the role of HDAC6i on metabolism in MM, we treated JJN3 and H929 MM cell lines with BAS-2 (HDAC6i) and measured glycolysis using extracellular acidification rate (ECAR) by seahorse. Mean basal glycolysis and glycolytic capacity were both reduced across three independent experiments following treatment with 3 μM BAS-2 (P = 0.024 and P =0.048, respectively; Fig. 1a,b). Additionally, we noted that HDAC6 is highly expressed across a panel of MM cell lines and HDAC6i causes an on-target increase in acetylated α-Tubulin, a HDAC6 substrate (Supplementary Fig. 1a,b). TTCP-10, a PROTAC our lab developed, caused an on-target degradation of HDAC6 (Supplementary Fig.1c). HDAC6i (BAS-2) and HDAC6 degradation (TTCP-10) caused a dose-dependent increase in cell death in a panel of MM cell lines (Supplementary Fig. 1d-e) and super-resolution STED live cell microscopy showed altered mitochondrial morphology following BAS-2 treatment. Subsequent Imaris surface rendering indicated a significant reduction in surface area per mitochondrion (Supplementary Fig. 1f), confirming that the compounds are working in a similar manner in MM, as we previously found in TNBC ^28,29^.

**Figure 1.**
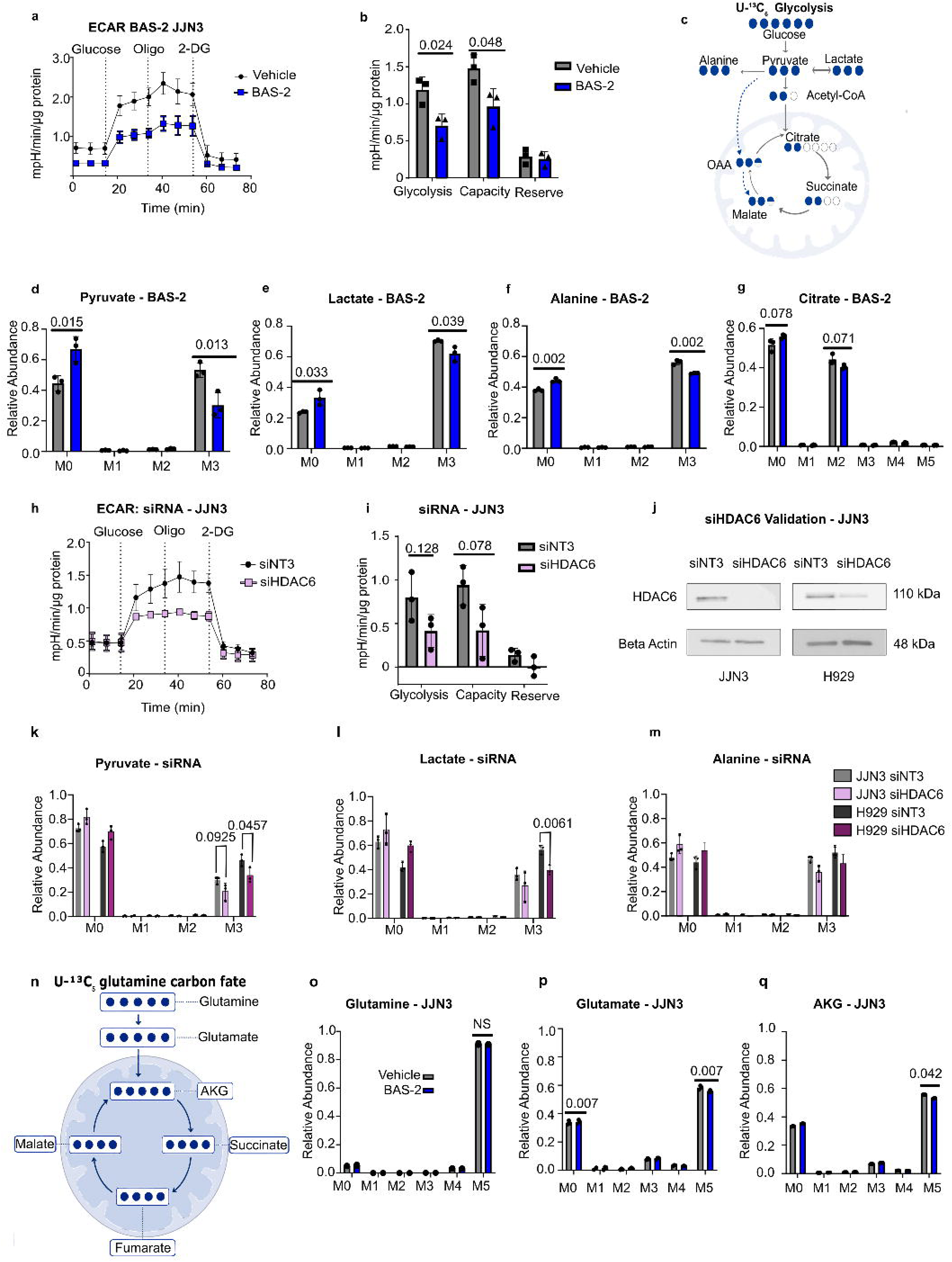
HDAC6 inhibition by BAS-2 or HDAC6 knockdown reduces glycolysis in MM. (**a**) Seahorse ECAR of JJN3 cells treated with DMSO versus 3 μM BAS-2, normalised to protein content per well (BCA assay). *n* = 3 independent biological replicates. (**b**) Glycolytic readouts derived from (**a**), DMSO versus 10 μM BAS-2. *n* = 3 independent biological replicates; unpaired two tailed t test with Welch correction. (**c**) Schematic of U-¹³C₆-glucose labelling. (**d–g**) Relative mass isotopologue abundance following a 30 minute U-^¹³C₆^glucose pulse in JJN3 MM cells treated with 10 μM BAS-2 versus DMSO. *n* = 3 independent cell cultures, analysed in a single GC–MS run; unpaired two tailed t test per isotopologue. (**h**) Seahorse extracellular acidification rate (ECAR) of JJN3 siHDAC6 versus siNT3 control cells, normalised to protein content per well (BCA assay). *n* = 3 independent biological replicates. (**i**) Glycolytic readouts derived from **(i)**. *n* = 3 independent biological replicates; unpaired two tailed t test. (**j**) Western blot validation of siRNA-mediated *HDAC6* knockdown in JJN3 and H929 MM cells. (**k–m**) Relative mass isotopologue abundance following a 30 minute U-^¹³C₆^ glucose pulse in H929 and JJN3 siHDAC6 versus siNT3 cells. *n* = 3 independent cell cultures, analysed in a single GC–MS run; unpaired two tailed t test per isotopologue. (**n**) Schematic of U-¹³C₅-glutamine tracing. (**o-q**) U-¹³C₅-glutamine 30 min pulse tracing in JJN3 MM cells, 10 μM BAS-2 versus DMSO. *n* = 3 independent cell cultures, analysed in a single GC–MS run; unpaired two tailed *t* test.

To identify effects on glycolytic flux, we performed 30-minute U-^13^C_6_-glucose pulse tracing in JJN3 cells treated with BAS-2 or vehicle and analysed mass isotopologue distributions of glycolytic intermediates by gas chromatography–mass spectrometry (GC-MS) (Fig 1c). BAS-2 reduced fully labelled M+3 pyruvate (P = 0.015, Fig. 1d), M+3 lactate (P = 0.033, Fig. 1e) and M+3 alanine (P = 0.002; Fig. 1f), with similar trends in M+2 citrate (P = 0.071, Fig. 1g), confirming that HDAC6i reduces glycolytic flux in MM. M+0 (unlabelled) pools were also higher in BAS-2 conditions, reflecting reduced contribution of glucose-derived carbon.

To confirm that the metabolic effects of BAS-2 reflect on-target HDAC6 inhibition, we depleted HDAC6 in JJN3 using siRNA and measured glycolysis using ECAR (Fig. 1h, i) and U-^13^C_6_-glucose pulse tracing. Knockdown of HDAC6 was confirmed by Western blot (Fig. 1j). Glucose-derived isotopologues of M+3 pyruvate (P= 0.049, Fig. 1k), M+3 lactate (P=0.006, Fig. 1l), M+3 alanine (P=0.044, Fig. 1m), showed reductions in the siHDAC6 compared to siNT3 in both the JJN3 and in the H929 cell lines (Supplementary Fig. 2a-c). The reductions were comparable to those seen after BAS-2 treatment indicating that loss of HDAC6 activity is responsible for the repression of glycolysis.

Breakdown of glutamine into glutamate and subsequently alpha-ketoglutarate (α-KG) is a common carbon-source pathway in many proliferating cells^8,10,30^. To test if glutaminolysis was being induced upon glycolysis repression, we performed a 30-minute U-^13^C_5_-glutamine pulse in JJN3 cells treated with 10 μM BAS-2 or vehicle for 24 h, and quantified isotopologue distributions of glutamine and its downstream metabolites by GC–MS (Fig. 1n) (Supplementary Fig. 2d-h). Interestingly, intracellular glutamine isotopologues were unchanged (Fig. 1o), and if anything, M+5 glutamate (P = 0.007, Fig. 1p) and α-KG (P=0.042, Fig. 1q) were reduced. These data indicate that BAS-2 does not redirect glutamine-derived carbon into the TCA cycle to compensate for blocks in glucose catabolism.

### 2.2 HDAC6i and depletion increase amino acid transporters and induce auxotrophy for amino acids

Previously, we performed data-independent acquisition (DIA) whole-cell proteomics on JJN3 cells treated with either HDAC6i (BAS-2) (Fig. 2a) or HDAC6-PROTAC (TTCP-10) (Fig. 2b; Supplemental Fig. 3a) relative to control, to demonstrate binding affinity^22^. Directional KEGG pathway enrichment indicated BAS-2 repressed metabolic pathway programs, including the TCA cycle, biosynthesis of amino acids and carbon metabolism, while enriching for alanine, asparagine and glutamine metabolism, along with endocytosis (Fig. 2c). To understand key proteins altered by both compounds, we performed protein-protein interaction analysis of differentially abundant proteins in both the TTCP-10 and BAS-2 treatments. Interestingly, this revealed coordinate upregulation of proteins related to amino acid import (Fig. 2d). Individual analysis of the four amino acid transporters indicated significant upregulation following HDAC6 inhibition or degradation, of SLC1A5, SLC7A5, SLC7A1 and SLC3A2 (Fig. 2e). These transporters cover the principal cellular routes of uptake in a variety of amino acids: SLC1A5 (ASCT2) imports asparagine, glutamine, serine and threonine; SLC7A5 (LAT1) heterodimerises with SLC3A2 (CD98) to import large neutral amino acids in exchange for intracellular glutamine; and SLC7A1 (CAT1) imports arginine and lysine^31–33^. The concerted upregulation of these proteins indicated that HDAC6-inhibited cells expand their capacity for extracellular amino-acid acquisition. A Western blot for SLC1A5 showed upregulation in JJN3 MM cells following both BAS-2 and TTCP-10 treatment (Supplementary Fig. 3b,c).

**Figure 2.**
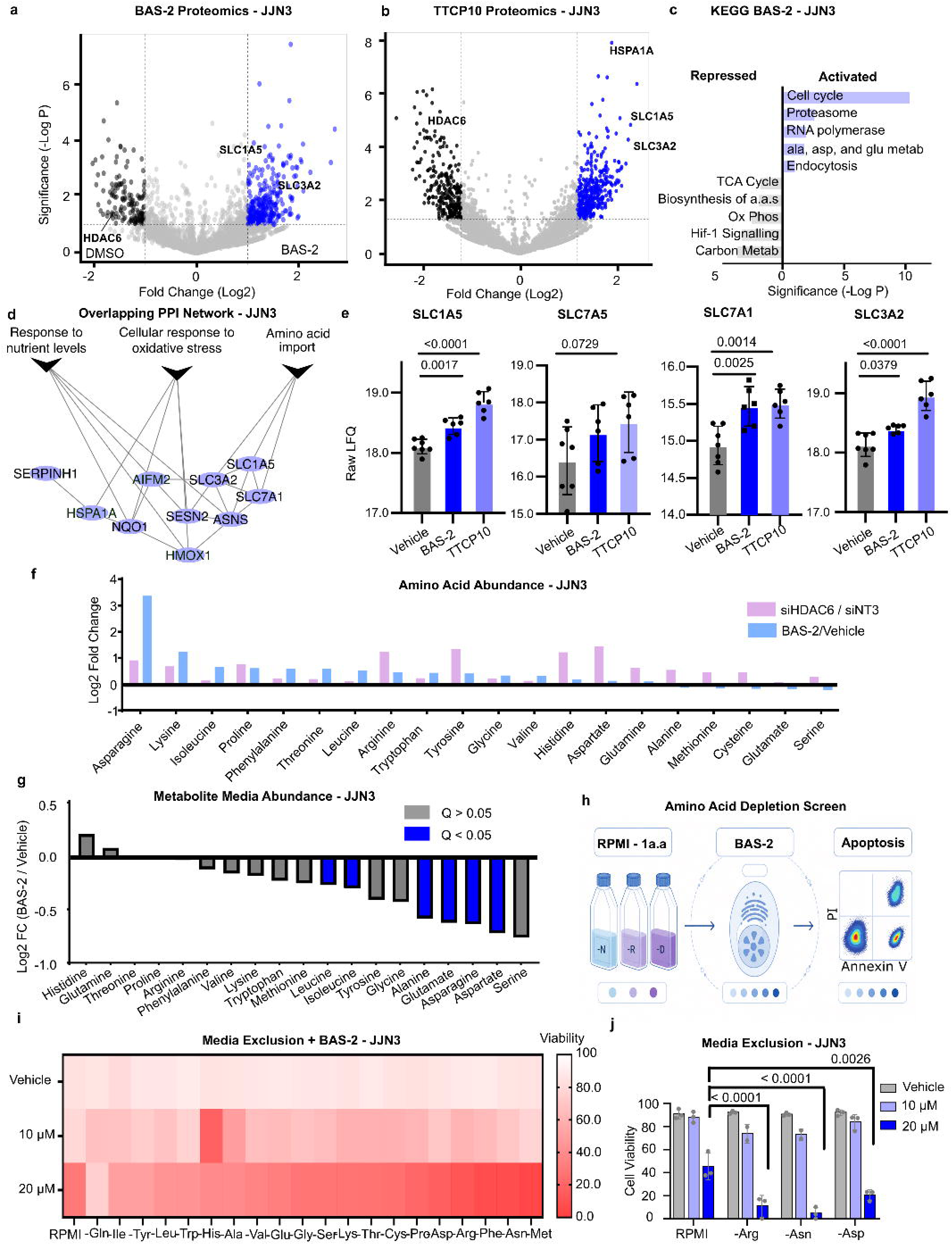
BAS-2 and HDAC6 knockdown remodel the intracellular amino acid pool. (**a**) Volcano plot of differentially abundant proteins by whole-cell proteomics, 10 μM BAS-2 versus DMSO, JJN3 MM cells. *n* = 3 independent biological replicates. Cutoffs P < 0.05 and FC > 1. (**b**) Volcano plot of differentially abundant proteins by whole-cell proteomics, 10 μM TTCP10 versus DMSO, JJN3 MM cells. *n* = 3 independent biological replicates. Cutoffs P < 0.05 and FC > 1. **(c**) KEGG enrichment analysis of activated and repressed metabolic pathways among cells treated with BAS-2. Proteins with P < 0.05 and FC > 1 were included. (**d**) Overlapped enrichment network of proteins upregulated in BAS-2 and TTCP10 Proteomics. (**e**) log₂-transformed LFQ intensities for key amino acid transporters, taken from whole-cell proteomics of JJN3 MM cells treated with DMSO versus 10 μM BAS-2. *n* = 3 independent biological replicates; unpaired two tailed *t* test. (**f**) Relative intracellular amino acid levels in siHDAC6 versus siNT3 cells and in DMSO versus 10 μM BAS-2 treated cells, shown as *Z* scores. *n* = 3 independent cell cultures, acquired within a single mass spectrometry batch normalized to cell number. (**g**) Extracellular amino acid concentrations shown as log₂ fold change (10 μM BAS-2 / DMSO). *n* = 3 independent cell cultures; multiple unpaired *t* tests with Benjamini–Yekutieli correction; significant *q* values indicated in blue, normalized to cell number. (**h**) Schematic of the amino acid depletion screen. (**i**) Heatmap of Annexin V–PI screen results following single amino acid depletion in JJN3 MM cells. Single exploratory screen (*n* = 1 biological replicate); no statistical testing was performed, and hits were validated independently in (**j**). (**j**) Validation of arginine, aspartate and asparagine depletion with 10 and 20 μM BAS-2 versus DMSO in JJN3 MM cells. Ordinary two-way ANOVA.

To investigate the intracellular abundance of amino acids, we analysed metabolite pools from the GC-MS experiments following treatment with BAS-2 or HDAC6 siRNA. Across the amino-acid panel, HDAC6 knockdown or BAS-2 treatment caused the accumulation of intracellular amino acids. In particular, asparagine was greatly increased intracellularly following BAS-2 treatment compared to control, however, there was elevations spanning the majority of measured amino acids relative to control (Fig. 2f); a result we previously observed in TNBC ^26^.

To confirm that elevated transporter expression, along with intracellular pools, translates into increased amino-acid uptake, we measured spent-medium amino-acid concentrations after 24 h of vehicle or 10 μM BAS-2 treatment of JJN3 cells, by targeted GC–MS. BAS-2-treated cells depleted extracellular amino acids relative to vehicle controls across the majority of the measured panel (Fig. 2g). Glutamine levels in the medium were slightly increased relative to control, consistent with its role as the intracellular exchange substrate for LAT1/CD98-mediated import of large neutral amino acids (Fig. 2g). Interestingly, while glutamate concentrations dropped in BAS-2-treated cells, we observed a depletion of glutamate from media, a finding that warrants further investigation. Of the 20 proteinogenic amino acids assessed, the top significant amino acids were alanine, glutamate, asparagine, and aspartate following multiple testing correction (Fig. 2g).

We next asked whether this expanded amino-acid uptake confers a reliance on particular extracellular amino acids that could be exploited therapeutically (Fig. 2h). We cultured JJN3 cells in RPMI formulations individually depleted of each of the 20 amino acids, combined with vehicle, 10 μM or 20 μM BAS-2, and assessed viability by Annexin V/PI flow cytometry (Fig. 2i). Multiple amino-acid withdrawals reduced viability under BAS-2; however, three hits, asparagine, arginine and aspartate, stood out because of their interconnected role in central carbon metabolism (Fig. 2j).

Asparagine and arginine both require aspartate as a precursor for *de novo* biosynthesis through asparagine synthetase (ASNS) and argininosuccinate synthetase (ASS1), respectively, and aspartate itself is produced by transamination of oxaloacetate (OAA)^34–37^. Validation of single amino-acid withdrawal of arginine, asparagine or aspartate confirmed significant reductions in cell viability (P < 0.0001, P < 0.0001 and P = 0.0026 respectively; Fig. 2j), along with reduced proliferation (Supplemental Fig. 3d-f), when treated with 20 μM BAS-2, establishing each as a BAS-2-induced amino-acid dependency.

Together, the elevated amino acid transporter expression, intracellular amino acid pools, the reduced amino acid in media, along with the enhanced cell killing following specific amino acid depletions, indicate that HDAC6-inhibited cells are increasing their propensity for amino acid uptake.

### 2.3 L-ASN treatment sensitises multiple myeloma cells to BAS-2

Having established that BAS-2 induces reliance on asparagine import in MM, next we wanted to test whether pharmacological depletion of asparagine using L-ASNase would synergise with BAS-2 in MM. L-ASNase hydrolyses extracellular asparagine to aspartate and to a lesser degree glutamine to glutamate, thus depleting asparagine and glutamine pools available to cells (Fig. 3a)^38,39^. We treated four MM cell lines (JJN3, H929, SKMM2, MM1S) with dose matrices of BAS-2 (variable 0–30 μM range tailored per cell line) and L-ASNase (0, 0.1, 1.0, 5.0 IU ml^-1^), with L-ASNase dosed on Day 1, followed by BAS-2 on Day 2 to capture the order in which the amino acid depletion screen was performed (Fig. 3b). Viability was quantified by Annexin V/PI flow cytometry at Day 3, and synergy was scored from the resulting matrices using the Loewe model in SynergyFinder^40^. All four lines exhibited positive synergy, with Loewe scores of JJN3 (16.3, P = 3.81 × 10^-3^), H929 (26.6, P = 1.40 × 10^-2^), SKMM2 (39.6, P = 1.54 × 10^-8^) and MM1S (19.1, P = 1.11 × 10^-3^)(Fig. 3c, Supplemental Fig. 4a-f).

**Figure 3.**
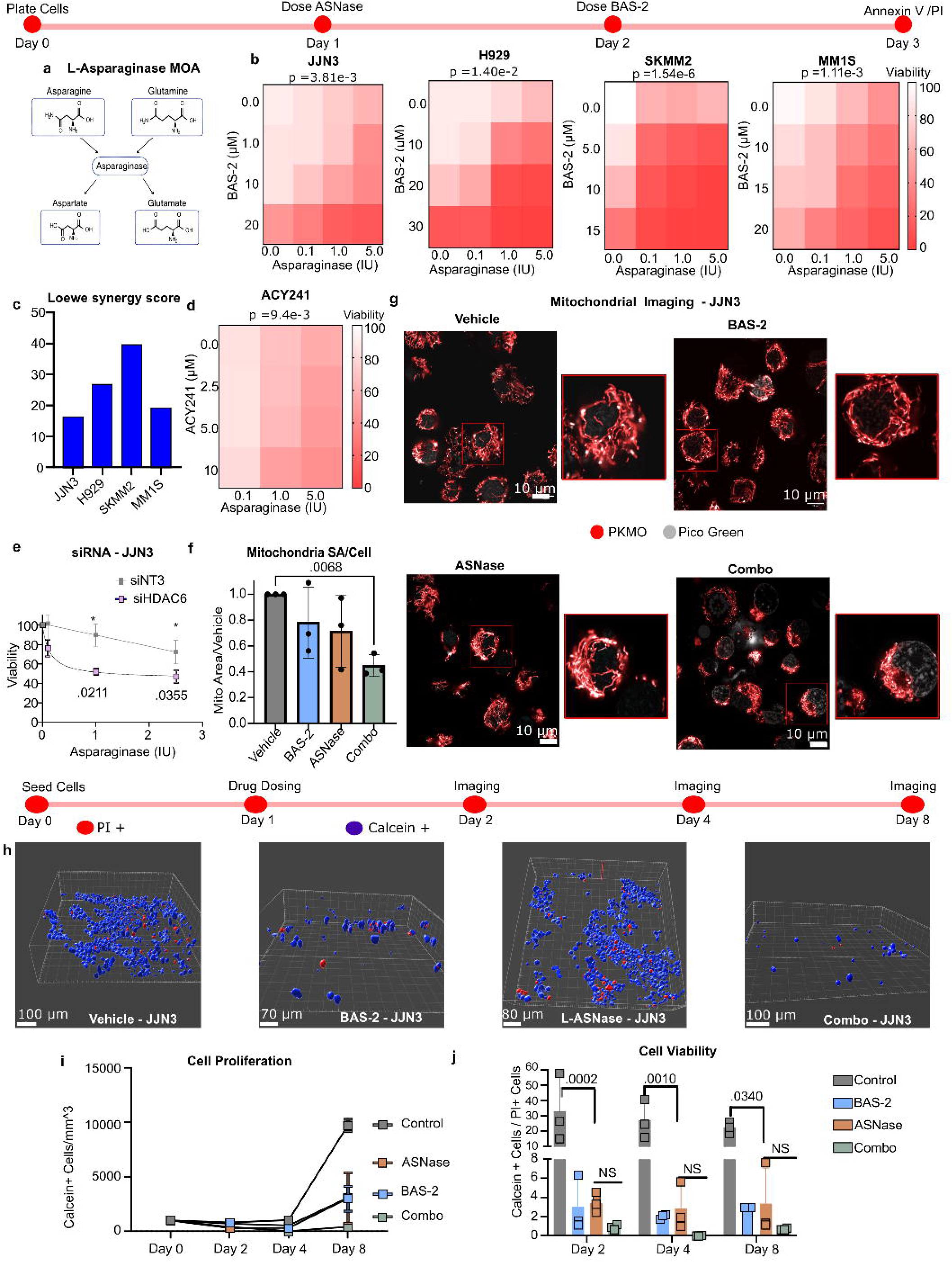
HDAC6 inhibition and depletion sensitise MM cells to L-asparaginase. (**a**) Schematic of the L-asparaginase (L-ASNase) mechanism of action. (**b**) Heatmaps of the cooperative cytotoxic effect of BAS-2 and L-ASNase by Annexin V–PI apoptosis screening. Data are mean cell viability across *n* = 3 independent biological replicates. P value was determined using SynergyFinder 3.0. (**c**) BAS-2 / L-ASNase Loewe synergy scores across four MM cell lines. Synergy calculated from *n* = 3 independent biological replicates per cell line via SynergyFinder 3.0. (**d**) Heatmap of the cooperative cytotoxic effect of ACY-241 and L-ASNase by Annexin V–PI apoptosis screening in JJN3 MM cells. Data are mean cell viability across *n* = 3 independent biological replicates. P value was determined using SynergyFinder 3.0. (**e**) Viability of JJN3 siHDAC6 versus siNT3 cells by CellTiter-Glo. *n* = 3 independent biological replicates; multiple unpaired t tests. (**f**) Imaris-reconstructed mitochondrial surface area relative to control. Total mitochondrial area per image was normalised to cell number per image, representing one technical replicate; the mean of all images per condition per experiment was taken as one biological replicate, cells sectioned by image border were excluded. Each data point represents an independent experiment; n = 3 biological replicates, one way ANOVA. (**g**) Representative STED images of mitochondria in JJN3 MM cells following BAS-2 (10 μM), ASNase (0.1 IU/mL), and combination treatment. n = 3 imaging experiments; scale = 10 μm. (**h**) Representative day-7 calcein/PI images of JJN3 MM cells embedded in RASTRUM 3D bioprinted matrices; surfaces reconstructed by Imaris rendering. Images acquired across 2 independent experiments. Scale bar length between 70 and 100 μm. (**i**) Relative number of calcein-positive cells over 7 days post rastrum seeding. Each data point represents one centrally acquired *z*-stack normalised to *z*-stack volume. Images acquired across 2 independent experiments; JJN3 MM cells. two-way ordinary ANOVA. (**j**) Ratio of calcein-positive to PI-positive cells over 7 days following RASTRUM seeding. Each data point represents one centrally acquired *z*-stack normalized to total cell density. Images acquired across 2 independent experiments; JJN3 MM cells. Two-way ordinary ANOVA.

To determine whether the synergy reflects a class HDAC6 inhibitor effect or one specific to BAS-2, we performed parallel L-ASNase dose matrices in JJN3 cells with two alternative HDAC6 inhibitors, ACY-241 (Fig. 3d) and the tool inhibitor tubacin (Supplementary Fig. 4j)^18,41^. We confirmed the enhanced cell killing of L-ASNase in siHDAC6 cells compared to siNT3 in both the JJN3 (Fig. 3e). We also assessed the combination in stable HDAC6 K/D JJN3 using CRISPR/Cas9 (sgHDAC6), where sgHDAC6 showed a slight increase in cytotoxicity to L-ASNase (Supplementary Fig. 4g,h).

To assess the mitochondria structure of the cells after single agent or combination treatment, we used PKMO live cell imaging by STED. Mitochondrial surfaces were reconstructed in Imaris and mitochondrial surface area per cell was quantified and normalized to cell number per image (Fig. 3g). Both BAS-2 and L-ASNase alone reduced per-cell mitochondrial surface area relative to vehicle, but the combination caused the greatest reduction (P = 0.0068) (Fig. 3f). To confirm the activity of the combination treatment in a more physiologically relevant setting, we seeded JJN3 cells in 3D bioprinted bone-marrow scaffolds using the Rastrum platform (Fig. 3h). Calcein-AM and propidium iodide co-staining were imaged at Days 2, 4, and 8, and Imaris surface reconstruction was used to quantify Calcein^+^ live cells (blue) and PI^+^ dead cells (red) within each organoid. Two readouts were extracted from these images: proliferation (Calcein^+^ cell number per mm^9^, normalised to organoid volume) and cell viability (Calcein^+^/PI^+^ ratio per region of interest). Both cell proliferation (Fig. 3i) and cell viability were reduced with the combination treatment, which was most obvious at day 8 (Fig. 3j). Lastly, to test whether the combination retains activity in the presence of bone-marrow stromal support, we repeated the Rastrum experiment with JJN3 cells co-printed with HS5 bone-marrow stromal cells tagged with mCherry (Supplementary Fig. 4i-k). Again, the combination retained its activity in a 3D bone-marrow-like environment, including under stromal co-culture, supporting translational relevance. In summary, HDAC6i or K/D enhanced L-ASNase activity across a panel of cell lines and in 3D bone-like environment.

### 2.4 L-ASNase enhances asparagine production from pyruvate

To define the metabolic state underlying combination-induced cell death, we profiled metabolite pools in JJN3 cells following 48 h treatment with 0.1 IU/mL L-ASNase. Amino acids dominated the dysregulated species in L-ASNase-treated versus control cells. As expected, asparagine and glutamine were the most significantly depleted species (P < 0.0001 and P = 0.0191, respectively) in the comparison (Fig. 4a), and targeted quantification confirmed significant depletion of both asparagine (Fig. 4b) and glutamine (Fig. 4c). To determine how asparaginase impacts central carbon metabolism, we performed a 24 h U-^13^C_6_-glucose tracing experiment, mapping glucose-derived carbon into glycolytic intermediates, the TCA cycle, and the downstream amino acids (Fig. 4d). Following BAS-2 (p= 0.009) and combination treatment (p= 0.0432), absolute M+3 pyruvate abundance decreases significantly (Fig. 4e). This drop implies a reduction in glucose-derived pyruvate for downstream carbon metabolic processes such as amino acid metabolism and TCA cycling following BAS-2 treatment. Interestingly, L-ASNase treatment increased the expression of the enzyme asparagine synthetase (ASNS) in whole-cell proteomics (p = 0.0098; Fig. 4f), with a goal of increasing *de-novo* asparagine synthesis from aspartate. The increased ASNS expression was paired with a sharp increase in absolute M+3 asparagine pools in the L-ASNase treated (p <0.001) group; *de-novo* asparagine synthesis drops in the combination treated cells by comparison (p = 0.0002, Fig. 4g).

**Figure 4.**
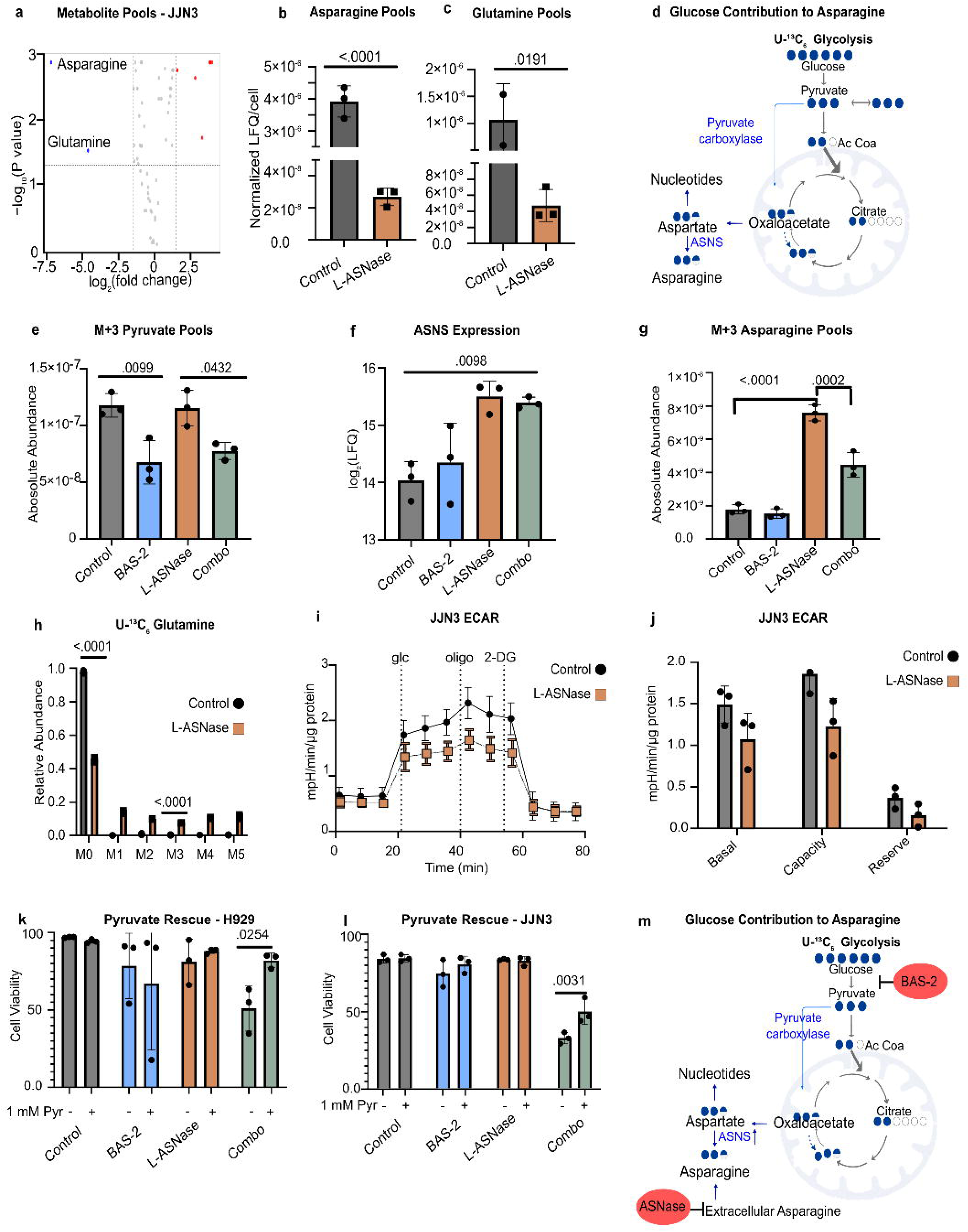
Combination treatment induces reliance on glucose catabolism paired with glycolytic block. (**a**) Volcano plot of differentially abundant metabolites following 0.1 IU/mL L-ASNase treatment in JJN3 MM cells; n = 3 independent cell cultures, analysed in a single GC–MS run. Multiple unpaired t test with Benjamini and Hochberg correction. (**b–c**) Asparagine and glutamine pools following 0.1 IU/mL L-ASNase treatment; n = 3 independent biological replicates; unpaired t test. (**d**) Schematic of U-¹³C₆-glucose tracing. (**e**) Absolute M+3 pyruvate abundance, calculated as the M+3 fractional enrichment multiplied by total sample pyruvate abundance, in JJN3 MM cells. *n* = 3 independent cell cultures, analysed in a single GC–MS run; one way ANOVA. (**f**) Asparagine synthetase (ASNS) abundance (log₂-transformed LFQ intensity) from whole-cell proteomics of JJN3 MM cells. *n* = 3 independent biological replicates; one way ANOVA. (**g**) Absolute M+3 asparagine abundance, calculated as the M+3 fractional enrichment multiplied by total sample asparagine abundance, in JJN3 MM cells. *n* = 3 independent cell cultures, analysed in a single GC–MS run; one way ANOVA. (**h**) Mass isotopologue distributions in JJN3 MM cells of glutamine, control versus 0.1 IU/mL L-ASNase. *n* = 3 independent cell cultures, analysed in a single GC–MS run; two-way ANOVA. (**i–j**) Seahorse extracellular acidification rate (ECAR) traces and the corresponding glycolytic readouts for JJN3 MM cells. *n* = 3 independent biological replicates, each the mean of technical replicates; two-way ANOVA. (**k–l**) Annexin V–PI analysis of cells cultured in RPMI with or without 1 mM exogenous pyruvate. *n* = 3 independent biological replicates; multiple unpaired *t* tests with uncorrected *P* values. JJN3 (**k**) and H929 (**l**) cell lines treated with 10 μM BAS-2, 1 IU/mL L-ASNase, and combo. (**m**) Proposed mechanism of synergy.

L-ASNase shifted the TCA cycle towards glucose-derived carbon: unlabelled M+0 fractions fell (P < 0.0001 for all four species) and higher isotopologues rose for glucose derived glutamine (Fig. 4h), succinate, fumarate and α-ketoglutarate (Supplementary Fig. 5d-i). Because glutamine is depleted under L-ASNase, we aimed to understand if this rise in fractional labelling was reflecting a loss of alternative carbon sources, or driven by an increase in glycolysis. We sought to understand if glucose catabolism increases as a compensatory mechanism following loss of glutamine contribution to carbon metabolism. Seahorse ECAR analysis showed that while BAS-2 reduces glycolysis in H929 and JJN3 cell lines, L-ASNase treatment did not increase basal glycolysis or glycolytic capacity relative to control (Fig. 4j, 4k) (Supplemental Fig. 5d-f; 5n-q).

Taken together, these data indicate that asparagine *de novo* synthesis is upregulated following asparagine depletion, and subsequent repression of glycolysis by BAS-2 restricts M+3 pyruvate and a capacity to convert to M+3 aspartate to M+3 asparagine. We reasoned that supplying carbon downstream of the BAS-2 glycolysis block should relieve this stress. Exogenous pyruvate rescued the viability of combination-treated H929 (Fig. 4k, p=0.0254) and JJN3 cells (Fig. 4l; p = 0.0031), supporting that the loss of glucose-derived carbon flow, rather than an off-target effect, drives the combination lethality (Fig. 4m).

### 2.5 Co-dosing reduces tumour burden in the chorioallantoic membrane model

To determine whether the metabolic mechanism characterised *in vitro* operates in a tumour setting, we engrafted JJN3 cells onto the chick chorioallantoic membrane and administered vehicle, 10 mg kg^-1^ BAS-2, 10 IU L-ASNase alone, or the combination at Day 3 and Day 5, harvesting tumours at Day 7 post implantation (Fig. 5a)^42^. To quantify anti-tumour *efficacy in ovo*, we measured tumour volume at Day 7 (EDD 16) across the CAM cohort. Combination-treated tumours showed the greatest reduction in tumour volume relative to vehicle controls, with BAS-2 and L-ASNase monotherapies producing intermediate reductions, at the doses utilized (Fig. 5b, p= 0.0068). To assess the malignant plasma cells specifically, we performed CD138^+^ immunohistochemistry on harvested tumours following fixation. The CD138-positive area was progressively reduced across monotherapy conditions and was lowest in combination-treated tumours (Fig. 5d, NS), indicating that the combination reduces CD138-positive plasma cell content *in vivo* beyond what either single agent achieves alone. As L-ASNase has known liver toxicity at higher doses ^43,44^, we also assessed the livers of the chicks following individual treatment or combination treatment. There were no obvious histological toxicities observed from the hematoxylin and eosin (H&E) staining in the liver (Fig. 5e) or in the, small intestine, brain, and heart (Supplementary Fig. 6a-c).

**Figure 5.**
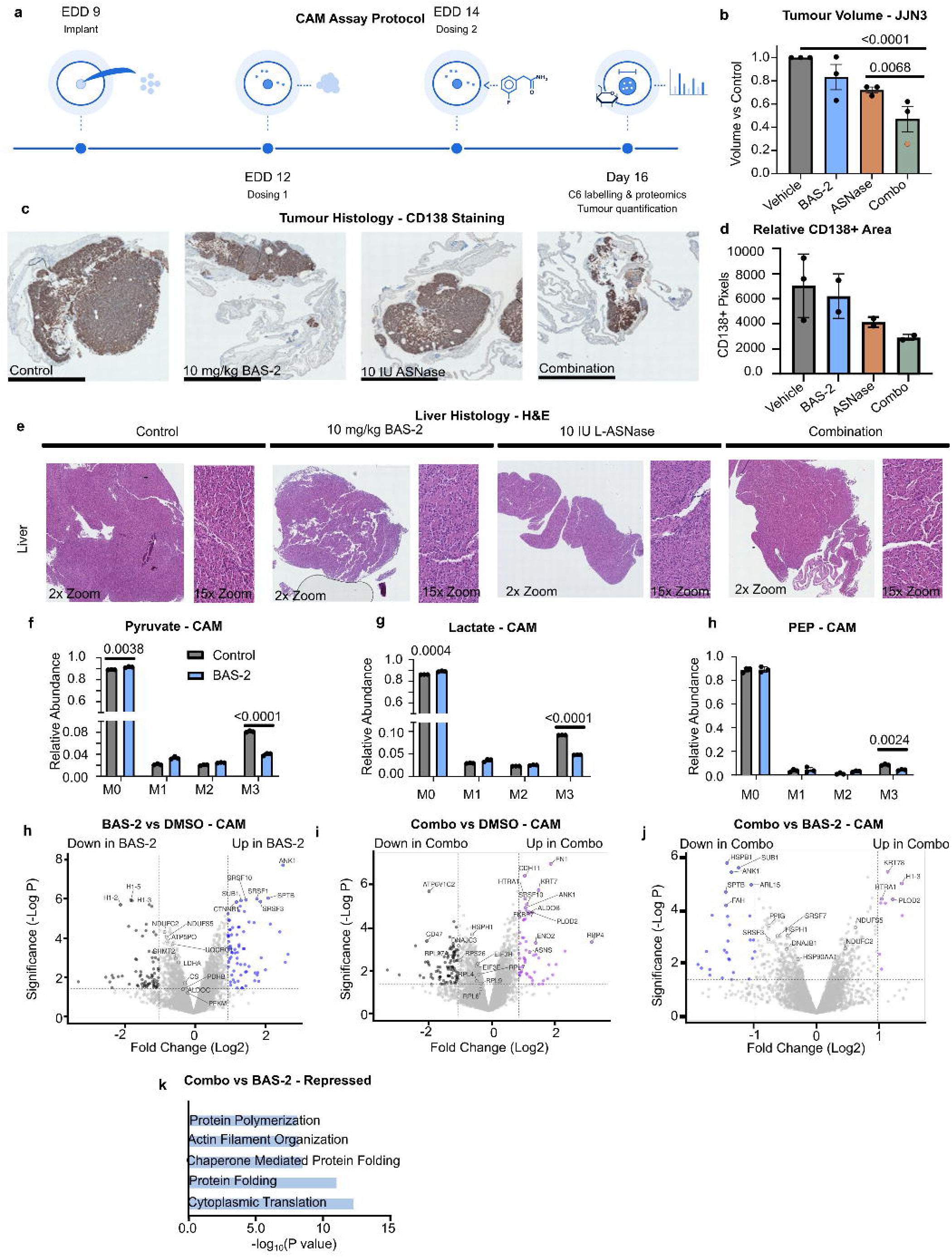
The BAS-2 / L-asparaginase combination is efficacious *in ovo* in the CAM model of MM. (**a**) Timeline of the chick chorioallantoic membrane (CAM) xenograft experiment. (**b**) Tumour volume relative to control, JJN3 MM xenograft. Each data point represents the mean of technical replicates. n = 3 independent biological replicates; one way ordinary ANOVA. (**c**) Representative CD138 tumour staining of resected tumours. *n* = 2 independent biological replicates (BAS-2, L-ASNase, Combo); n = 3 independent biological replicates (vehicle). Scale bar = 1 mm. (**d**) Quantification of CD138-positive staining. *n* = 2 independent biological replicates (BAS-2, L-ASNase, Combo); n = 3 independent biological replicates (vehicle). (**e**) Representative tumour H&E imaging of CAM resected organs from single arm treatments and co treated samples. (**f–h**) Relative mass isotopologue abundance following a 16-hour U-^¹³C₆-^glucose pulse in CAM tumour resections treated with 10 mg/kg BAS-2 versus DMSO. *n* = 3 tumours, analysed in a single GC–MS run; unpaired two tailed t test per isotopologue. (**h-j**) Volcano Plots of differentially regulated proteins, n = 3 resected tumours, CAM. Log₂ fold change cutoff = 1 and −log p value = 1.3. (**k**) Repressed pathways in combination-treated tumour resections relative to control. *n* = 3 independent biological replicates. GO enrichment. P value < 0.1 cutoff.

To confirm the molecular metabolic mechanisms identified *in vitro* were occurring *in vivo* we administered 24 h U-^13^C_6_-glucose to a separate cohort of BAS-2 and vehicle treated CAM tumors and quantified pyruvate isotopologues by GC-MS. BAS-2-treated tumours showed a clear reduction in M+3 pyruvate relative to vehicle controls (Fig. 5f, p <0.0001), as well as M+3 lactate (Fig. 5g, p <0.0001) and PEP (Fig. 5h, p=0.0024), as well as other expected metabolic signatures (Supplemental Fig. 6d-h), confirming that BAS-2 inhibits glycolysis *in ovo* in an MM xenograft. We additionally analysed metabolite abundance from resected liver species of chicks treated with BAS-2, ASNase, and combination. Our findings suggest drops in majority of metabolites following both BAS-2 and ASNase treatment (Supplementary Fig. 6j)^43,44^.

To determine the signalling induced by the combination treatment *in ovo*, we performed whole-tissue proteomics on CAM tumours from all four treatment groups. Comparison of combination-treated tumours to vehicle controls produced distinct proteomic signatures (Fig. 5h-k). We performed bidirectional pathway enrichment analysis on differentially abundant proteins to define upregulated and downregulated processes in combination treated tumours versus BAS-2 treated (Supplemental Fig. 7a-k). Downregulated terms by GO enrichment were dominated by pathways related to mRNA processing, protein polymerisation, and protein folding^45,46^ (Fig. 5K), processes consistent with broad suppression of anabolic and translational programmes.

### 2.6 Co-dosing triggers integrated stress response and loss of viability in a primary sample

Given the protein pathways reduced by combination treatment, we next investigated the effects of combination treatment on new protein synthesis in living cells using O-propargyl-puromycin (OPP) incorporation in JJN3 cells. OPP is an amino acid analogue that can be fluorescently labelled to accurately measure protein synthesis rates by flow cytometry^47^. Control and BAS-2 treated cells maintained similar levels of protein synthesis, while rates were mildly reduced in L-ASNase-treated cells. However, the combination treatment strongly restricted protein synthesis relative to single arm L-ASN and control (Fig. 6a-b, P < 0.002 for both comparisons). This result supports that combination treatment reduces asparagine pools and flux sufficiently to impair translational capacity. The loss of protein synthesis under combination treatment, but not L-ASNase alone, suggests that glucose-derived carbon is maintaining asparagine pools and supporting translation during L-ASNase treatment, the addition of BAS-2 reduced pyruvate preventing protein translation.

**Figure 6.**
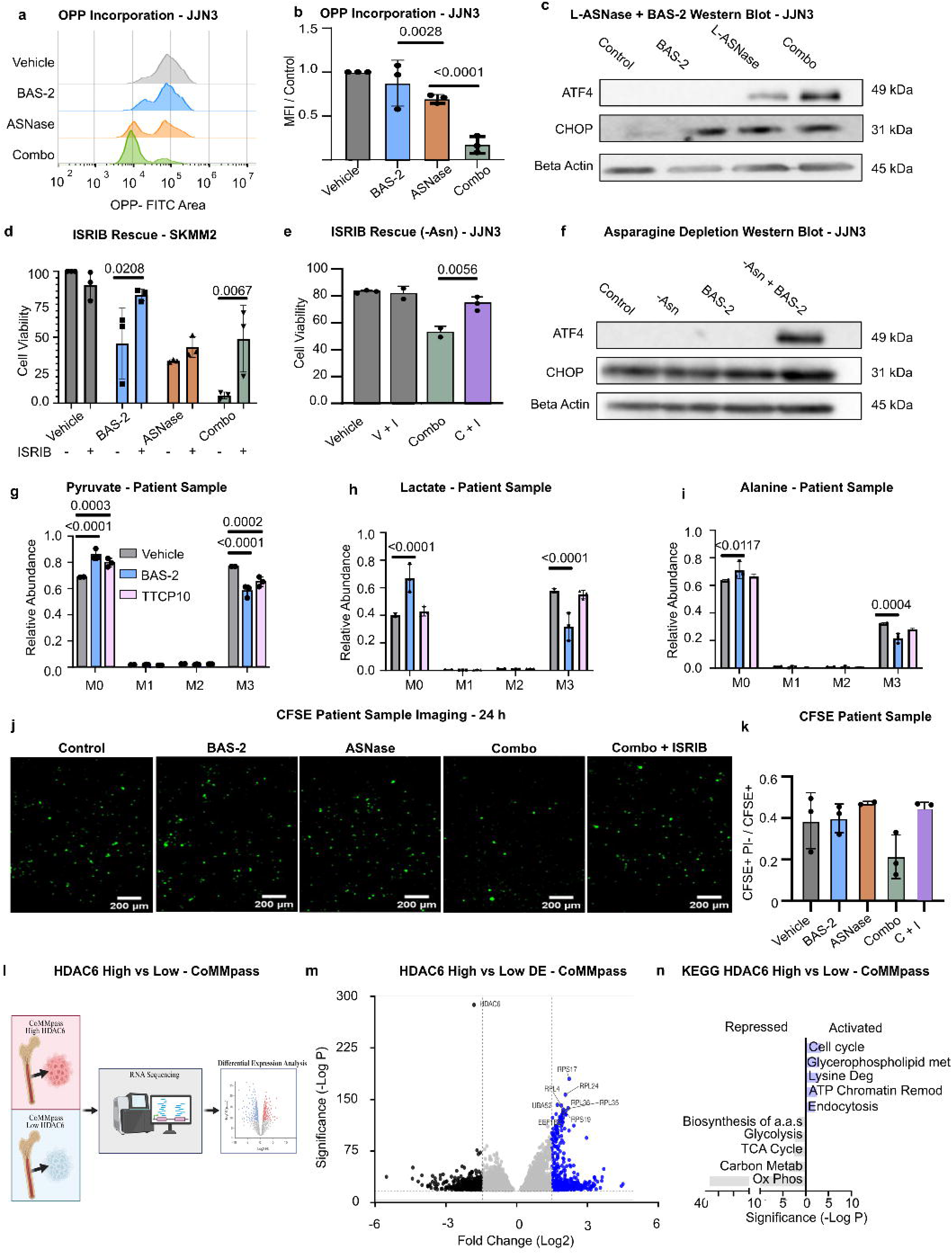
The BAS-2 / L-asparaginase combination activates the integrated stress response. (**a-b**) O-propargyl-puromycin (OPP) incorporation assay in JJN3 MM cells. *n* = 3 independent biological replicates; one-way ANOVA. (**c**) Western blot of JJN3 MM cells treated with BAS-2 and L-ASNase, probed ATF4 and CHOP, relative to Beta actin loading control. (**d**) Annexin V–PI of BAS-2 / L-ASNase co dosing in SKMM2 MM cell lines. Cells treated with 500 nM ISRIB, analysed via one way ANOVA. (**e**) Western blot of JJN3 cells treated with Control, BAS-2, L-ASNase or comb and proved for ATF4, CHOP and Beta actin loading control. **(f)** Western blot of JJN3 MM cells cultured in -ASN media with BAS-2, probed ATF4 and CHOP, relative to Beta actin loading control. (**g-i**) Relative mass isotopologue abundance following a 16 hour U-^¹³C₆-^glucose pulse in CD138+ primary tumour cells treated with 10 μM BAS-2, TTCP10 versus DMSO. *n* = 3 separate cultures of cells from one patient (technical replicates), analysed in a single GC–MS run; unpaired two tailed t test per isotopologue. (**j**) Maximum intensity projection of CFSE signal from bioprinted matrices 24 hours after implantation and treatment; all samples co-stained with PI. *n* = 2–3 bioprinted matrices, 200 μm scale bar. (**k**) Relative area of CFSE-positive, PI-negative signal as a proportion of total CFSE-positive signal per matrix. n = 3 independent scaffolds per group (control, BAS-2, L-ASNase, combo, combo + 100 nM ISRIB). (**l**), Schematic of the CoMMpass (IA14) analysis. The 100 transcriptomes with the highest and the 100 with the lowest *HDAC6* expression underwent differential expression analysis. (**m**), Volcano plot of differentially expressed genes between HDAC6-high and HDAC6-low tumours. Dashed lines mark the fold-change and significance thresholds (|log₂FC| > 1.5; adjusted *P* < 0.05). (**n**), KEGG pathways enriched among genes upregulated (activated) or downregulated (repressed) in HDAC6-high relative to HDAC6-low tumours.

Given the stark drop in protein synthesis, we postulated that global translation may be halted, potentially indicating triggering the integrated stress response (ISR). To confirm ISR activation biochemically, we performed Western blotting on CAM tumour lysates across all four treatment conditions. First, we confirmed acetylation of α-tubulin following BAS-2’s treatment *in ovo*, confirming HDAC6i. Additionally, phosphorylated eIF2α, a critical marker of ISR activation, and CHOP, the downstream cell death factor regulated by chronic ISR activation, were probed (Supplementary Fig. 8a).

We confirmed an induction of ATF4 and CHOP in either L-ASNase or combo treatment in JJN3 cells cultured *in vitro (*Fig. 6c,f). Addition of ISRIB, the inhibitor of ISR which stabilises eIF2B^48,49^ and restores translation downstream of eIF2α-P, reduced ATF4 induction (Supplemental Fig. 8b) and reduced cell death of the combination treatment in both SKMM2 (Fig. 6d, P = 0.0067) and JJN3 (Fig. 6e, P=0.0056, Supplementary Fig. 8c). These data indicate that limited asparagine leads to uncharged tRNA, global translation inhibition as measured by OPP and activation of ISR that is important for cell death and demonstrated by ISRIB, reducing cell death.

We were fortunate to attain a primary patient MM sample that had sufficient material for U-^13^C_6_-glucose tracing. We were able to confirm, that HDAC6i with BAS-2 or degradation with TTCP-10 also reduced glycolysis in primary MM sample. We incubated the cells with U-^13^C_6_-glucose for 16 hours and traced the labelled carbon to M+3 pyruvate (Fig. 6i, P=<0.0001), M+3 lactate (Fig. 6j, P=<0.0001) and M+3 alanine (Fig. 6j P=0.0117) post treatment, demonstrating a reduction of glycolysis in primary MM sample *ex-vivo* (Supplementary Fig. 8d-g). We also assessed the activity of the combination treatment of BAS-2 and L-ASNase in a separate MM patient sample grown in 3D using the Rastrum. CD138^+^ cells were stained with CFSE cell tracker and seeded in the bone-like matrigel. Following 24 hours of treatment, cells were co-stained with PI and imaged via whole-scaffold z-stack (Fig. 6j). Relative maximum projections of CFSE + PI-cells, which were used as viable cell readouts, showed a decreased density in combination treated cells that was restored to control levels following inhibition of the ISR. Quantification indicated a drop in the ratio of CFSE+ PI-objects over total CFSE+ objects indicating a shift in viability that was restored by ISR inhibition (Fig. 6k).

Lastly, to assess the role of HDAC6 in primary MM transcriptomes from the MMRF CoMMpass Ia14a dataset, we stratified the transcriptomes by HDAC6 expression^50^. Next, we assessed if HDAC6 co-varies with metabolic gene programs. Pathways enriched in the HDAC6-low group recapitulated those repressed by BAS-2, including glycolysis, TCA cycle, amino-acid biosynthesis (Fig. 6n). The two groups were broadly comparable across age, ECOG performance status, treatment class, transplant eligibility, and ISS/R-ISS stage (Supplementary Fig. 9).

## Discussion

Metabolism is notoriously difficult to target therapeutically, this is due to the inherent flexibility within the system. The current metabolic dependencies that are targetable arise from either a mutation driving an enzyme to produce an alternative metabolite, like IDH2 mutations or due to the lack of an enzyme like ASNS inducing a dependency on extracellular asparagine for protein ^9,15,16,51,52^. In this study, we showed that BAS-2 represses glycolytic flux in MM cells, this drives a reliance on import of amino acids, in particular asparagine, to maintain the high biosynthetic requirements of MM cells and exposes a metabolic vulnerability that can be targeted with L-ASNase.

Metabolic reprogramming is increasingly recognised as a driver of adaptive resistance^53,54^, yet few metabolic vulnerabilities in myeloma have proved tractable in the clinic. Our work addresses this gap by showing that an enzyme already of therapeutic interest in myeloma^24,55^, HDAC6, acts as a node of carbon metabolism, and that inhibiting it does not simply blunt metabolism but reroutes it, creating a new, targetable dependency with L-ASNase. MM arises in the plasma cell, a terminally differentiated B-cell. Prior studies have shown that asparagine import supports B-cell homeostasis^9,56^. B-cell acute lymphoblastic leukaemia lacks asparagine synthase, and thus lack the capacity for asparagine *de novo* synthesis, and are particularly sensitive to asparagine depletion^52,57–59^. Indeed, there is evidence of hypermethylation of asparagine synthetase in myeloma, suggesting a vulnerability to L-ASNase treatment^60^.

Of the amino acids analysed, asparagine showed the largest increase intracellularly following BAS-2 treatment. Asparagine has been described as an amino acid exchange factor, with intracellular levels governing the uptake of other amino acids^61^. HDAC6 inhibition raised intracellular asparagine alongside a coordinated upregulation of amino acid transporters, increased depletion of extracellular amino acids, and our depletion screen identified arginine, aspartate and asparagine. The convergent collapse of intracellular asparagine pools due to a reduction in pyruvate from BAS-2 and L-ASN depleting extracellular asparagine triggered the integrated stress response (ISR) activation and apoptotic cell death, an outcome rescued by exogenous pyruvate and by pharmacological ISR inhibition with ISRIB.

Previous reports have shown that loss of asparagine leads to activation of the ISR, sensed by general control nonderepressible 2 (GCN2) causing increased activating transcription factor 4 (ATF4) expression^60,62,63^. The ISR is a conserved pathway that allows cells to adapt to nutrient limitation and other stressors^64–69^. We have not directly identified the upstream sensor kinase responsible for eIF2α phosphorylation in this context, but the dependence of the response on intracellular asparagine availability is consistent with engagement of GCN2, which senses uncharged tRNAs and is the canonical sensor of amino-acid limitation^67,69^

A recent study in Nature Metabolism demonstrated that in cells with respiration deficiency, induced through mtDNA mutations at varying heteroplasmy, developed an auxotrophy for asparagine^70^. This study, identified a similar mechanism to our work, but with an opposing goal to rescue the cells, they showed that asparagine supplementation could protect the proliferation rate of respiration defective cells through blocking GCN2 activation, and preventing ATF4 activation.

Our data showed that the combination of HDAC6i and L-ASNase produces synergistic killing across a panel of myeloma cell lines, suppresses growth in 3D bioprinted bone marrow organoids from primary patient samples and cell lines, and reduces tumour burden in CAM xenografts. However, whether the combination is specific for MM, that have a high biosynthetic demand or whether it can be expanded to other tumour types is something we are currently investigating. Previously, we showed in TNBC cell lines that HDAC6 interacted with fumarate hydratase and altered fumarate levels, leading to an increase in the succination of proteins^26^. In MM cell lines, upon glucose labelling we did not detect major changes in fumarate levels, this could be due to a context-specific role for HDAC6 in TNBC, which have high FH amplification rate^28^. Interestingly, it was shown previously in TNBC that ASNS expression correlated with the potential to metastasise^71^. In that study, L-ASNase or dietary restriction of asparagine reduced the capacity to metastasise, without altering the primary tumour.

Clinically approved L-ASNases are exclusively of bacterial origin which renders them prone to immunogenic responses, necessitating strategies such as PEGylation which has only partially mitigated hypersensitivity reactions. Dose limiting toxicity is a prominent clinical issue in adult ALL patients, however recent progress, notably the development of guinea pig L-ASNase, may overcome immunogenicity due to its close sequence homology with L-ASNase, as well as fewer side effects, owing to its inherent glutaminase-free nature. This may make the use of this therapy more acceptable in the MM patient population^72^. One limitation of this work is that the CAM xenograft model permits rapid evaluation of efficacy and on-tumour mechanistic readouts *in vivo*, but it lacks the host immune competence and the long-term complexity of a mouse model. Despite these limitations, the central translational implication is clear: pan-HDAC inhibitors are already approved for MM, L-ASNase is approved for B-ALL, and the combination of selective HDAC6 inhibition with pharmacological asparagine depletion exploits a metabolic vulnerability defined by stable isotope tracing and validated in 3D and *in vivo* models^2,15^.

The convergence between drug-induced, HDAC6 gene silencing, and CoMMpass-patient-level HDAC6 metabolic signatures suggests that the dependencies revealed by BAS-2 reflect genuine HDAC6 biology in patient tumours and cytogenetic and survival differences in HDAC6-high/low cohorts may suggest expression of HDAC6 as a predictor of disease progression and type. This combination represents a clinically tractable, mechanistically grounded therapeutic approach for MM, and motivates continued preclinical development of selective HDAC6 inhibitors in combination with established asparagine-depleting therapies.

## Methods

### 5.1 Drug compounds and treatment scheme

BAS-2 (selective HDAC6 inhibitor) was synthesised in-house and dissolved in dimethyl sulfoxide (DMSO) to a desired stock concentration (1000x working dosage) immediately before use in each experiment. L-ASNase (Erwinase) was reconstituted in sterile PBS to a stock concentration of 2000 IU/ml and stored at -20 °C. ACY-241 (citarinostat) and Tubacin were obtained commercially, dissolved in DMSO to 10 mM stocks, and stored at -80 °C. ISRIB was reconstituted in DMSO to a 1 mM stock and stored at -20 °C in single-use aliquots. Sodium pyruvate (Sigma) was prepared at a 100 mM stock concentration in PBS and used at a final concentration of 1 mM in rescue experiments. For all drug treatments, final DMSO concentration in cell culture media was maintained below 0.1%.

For combination experiments, the standard treatment scheme was as follows: cells were plated on Day 0; L-ASNase was added on Day 1; BAS-2 was added on Day 2; and cells were harvested for downstream analysis on Day 3. Single-agent treatments followed the same timeline, with the omitted compound replaced by an equivalent vehicle. Unless otherwise specified, BAS-2 was used at 10 μM, which is non-cytotoxic as a monotherapy at this concentration in MM cell lines.

### 5.2 Cell Lines

JJN3, H929, MM1S, SKMM2, KMS12-PE, KMS27, KMS12BM, U266, RPMI8226, KMS18 and HEK293T cells were obtained from the American Type Culture Collection (ATCC). RPMI-1640 medium (Sigma-Aldrich) was used for all multiple myeloma cell lines and DMEM/F12 (Sigma-Aldrich) for HEK293T cells. All media were supplemented with 10% foetal bovine serum (FBS), 5 mg/ml penicillin/streptomycin, and 10 mM L-glutamine. Cells were maintained at 37 °C with 5% CO_2_ and split at 70% confluency.

Amino acid depletion screening was performed in JJN3 cells to validate findings of whole-cell proteomics. Hits from the screen were validated in H929 and MM1S cells. HEK293T cells were used for lentiviral particle production. SKMM2 was added as a fourth myeloma cell line for combination synergy analyses.

### 5.3 Western blotting

Cells or tumour tissue were lysed in RIPA buffer (Thermo Fisher, supplemented with protease and phosphatase inhibitors; Sigma-Aldrich) and quantified by BCA assay (Thermo Fisher). Samples were denatured at 95 °C in 1× Laemmli loading buffer. 10-20 μg of protein was loaded onto 10% polyacrylamide gels, separated by electrophoresis, and transferred to PVDF membranes (18 h transfer at 20 V @ 4°C). Membranes were blocked with 5% milk in tris-buffered saline with Tween-20 (TBS-T) for 1 h at room temperature. Primary antibodies were incubated overnight at 4 °C in 5% milk-TBS-T at 1:1,000 dilution. Horseradish peroxidase (HRP)-conjugated goat anti-rabbit or anti-mouse secondary antibodies were incubated for 1 h at room temperature before imaging via enhanced chemiluminescent substrate (Millipore) on a LAS-4000 imaging system (Fujifilm). Loading control antibodies were incubated afterwards on the same membrane. Densitometry was performed using ImageJ.

### 5.4 Antibodies

Primary antibodies included EIF2α (Cell Signaling Technology, 5324S), phospho-EIF2α Ser51 (Cell Signaling Technology, 9721L), ATF4 (Proteintech, 10835-1-AP), CHOP (Cell Signaling Technology, 2895S), HDAC6 (Cell Signaling Technology, 7558S), and acetyl-α-tubulin (Cell Signaling Technology, 5335S). β-actin was detected using an anti-actin antibody from Sigma-Aldrich (A5441). Secondary antibodies were obtained from Millipore, including anti-mouse (AP124P) and anti-rabbit IgG (AP132P). All primary antibodies were IgG isotypes unless otherwise specified.

### 5.26 Statistical analysis

All statistical analyses were performed in GraphPad Prism (v10) using default settings. Two-tailed tests were used throughout. For comparisons of two groups with a single independent variable, unpaired Student’s t-tests were applied. For comparisons of three or more groups with a single independent variable, one-way ANOVA was applied. For experiments with two independent variables (for example, drug × amino-acid exclusion, drug × ISRIB, or drug × exogenous pyruvate), two-way ANOVA was. For ^13^C isotopologue panels, one unpaired t-test was performed per isotopologue. For four-condition ^13^C isotopologue panels (Figure xx), two-way ANOVA was applied. Pool size comparisons in were analysed by one-way ANOVA. Significance is reported as exact P-values where space permits, otherwise as *P < 0.05, **P < 0.01, ***P < 0.001, ****P < 0.0001. Multiple testing corrections were used as specified in figure legends. Bars represent the mean and error bars represent the standard error of the mean (SEM) unless otherwise indicated. n refers to independent biological replicates, except for GC–MS-based tracing and metabolomics, where n refers to separate cell cultures analysed in a single GC–MS run (see Stable isotope tracing). Figures were prepared in Inkscape; schematics were generated in inkscape and BioRender.

## Data availability statement

The mass spectrometry proteomics data have been deposited in the ProteomeXchange Consortium via the PRIDE partner repository with the dataset identifier for BAS-2 and TTCP-10: PXD063860. The mass spectrometry proteomics data for the CAM JJN3 have been deposited to the ProteomeXchange Consortium via the PRIDE partner repository with the dataset identifier PXD087843 and 10.6019/PXD087843.

## Author Contributions

MH performed experiments, analysed data and wrote the paper. TNC conceived the project, designed experiments and wrote the paper. CD, IC, JF, KF, YW, SG, LDM, and AR, performed experiments or analysed data. NR and KW analysed the Mass spectrometry data. DAR, CD generated reagents for the paper. HD, MG, KW, NR, TS, and MW provided technical support and facility access. JQ and SVG provided patient samples. MW analysed metabolomics data and DAR, NR, HD, MG, MW, EK, CD, MM, AMH, JHMP, SVG and JQ edited the manuscript.

## Acknowledgements

The authors would like to thank the following funding agencies for their support: Fulbright supported MH; Irish Research Council Laureate Award (IRCLA/2022/2822) supported TNC, SG, LdM and CD. This work was supported TNC under Taighde Éireann-Research Ireland and Myeloma Ireland under POI-02 (23/SPP/12064). Confocal and STED images were acquired in the RCSI Super Resolution Imaging Consortium (SRIC) funded by Science Foundation Ireland (18/RI/5723). The Comprehensive Molecular Analytical Platform (CMAP) under The SFI Research Programme, reference 18/RI/5702, supported mass spectrometry research reported in this publication. Supported by funding for equipment 18/RI/5792 (Research Ireland) setup within the Molecular and Cellular Imaging Core of the Centre of systems Medicine, Dept. of Physiology and Medical Physics, RCSI.

## Notes

### Competing Interest Statement

The authors have declared no competing interest.

