## Supplementary Figures and Legends for "HDAC6 inhibition rewires carbon metabolism to expose an asparagine dependency in multiple myeloma"

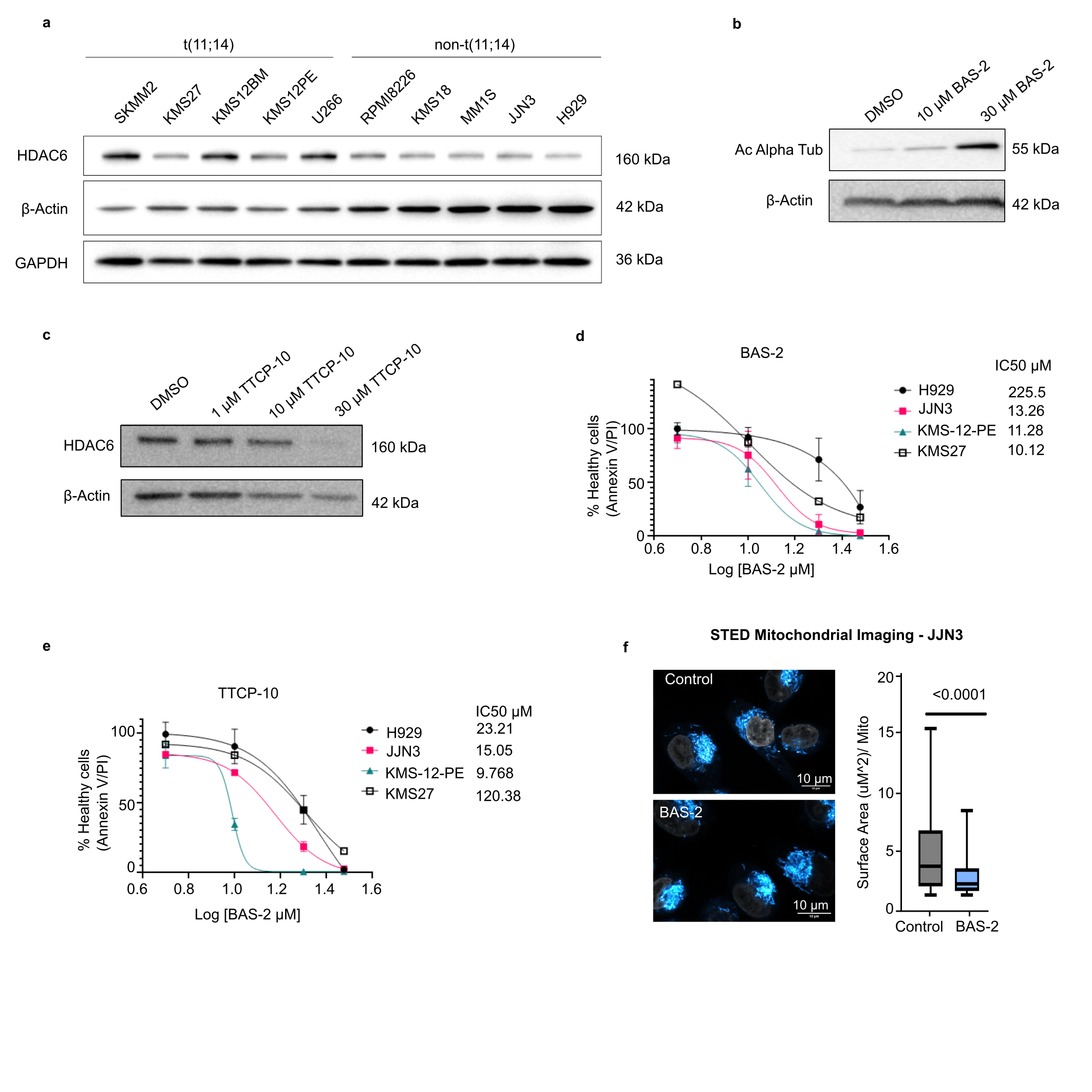

**Supplementary Fig. 1 | HDAC6 expression and activity of BAS-2 and TTCP-10 in MM cell lines.**

**(a)** Immunoblot of HDAC6 in t(11;14) (SKMM2, KMS27, KMS12-BM, KMS12-PE, U266) and non-t(11;14) (RPMI-8226, KMS18, MM1S, JJN3, H929) myeloma lines. β-actin and GAPDH are shown as loading controls. Representative of n = 3 independent blots. **(b)** Immunoblot of acetylated α-tubulin in JJN3 cells treated with DMSO or BAS-2 (10 or 30 μM) for 24 h; β-actin loading control. **(c)** Immunoblot of HDAC6 in JJN3 cells treated with DMSO or TTCP-10 (1, 10 or 30 μM) for 24 h; β-actin loading control. **(d,e)** Annexin V–PI dose–response curves for BAS-2 (d) and TTCP-10 (e) in H929, JJN3, KMS-12-PE and KMS27 cells after 24 h. Values are the percentage of Annexin V⁻/PI⁻ cells normalised to DMSO and plotted against log₁₀ dose (μM). IC₅₀ values (μM) are listed beside each line. Data are mean ± s.e.m.; n = 3 independent experiments. **(f)** Representative STED images of JJN3 mitochondria after vehicle or 10 μM BAS-2 for 24 h (left), with mitochondrial surface area quantified per mitochondrion (right). Box plots show median and interquartile range, with whiskers at minimum and maximum; unpaired two-tailed Student’s t-test. Scale bars, 10 μm.

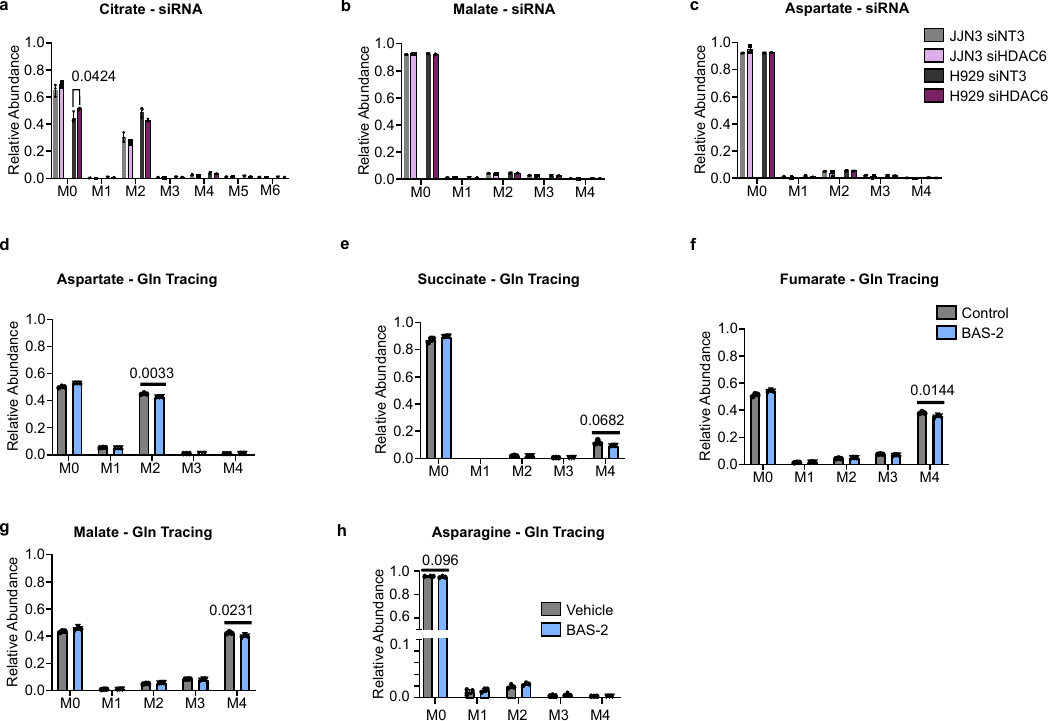

**Supplementary Fig. 2 | Isotope tracing in HDAC6-silenced cells and glutamine tracing.**

**(a–c)** U-¹³C₆-glucose isotopologue distributions of citrate (a), malate (b) and aspartate (c) in JJN3 and H929 cells transfected with non-targeting (siNT3) or HDAC6 siRNA (siHDAC6) and labelled for 30 min. n = 3 independent cell cultures, analysed in a single GC–MS run. Within each cell line, one unpaired two-tailed t-test was run per isotopologue. **(d–h)** U-¹³C₅-glutamine isotopologue distributions of aspartate (**d**), succinate (**e**), fumarate (**f**), malate (**g**) and asparagine (**h**) in JJN3 cells treated with vehicle or 10 μM BAS-2 for 24 h and labelled for 30 min. n = 3 independent cell cultures, analysed in a single GC–MS run; tests as in a–c. n = 3 independent cell cultures, analysed in a single GC–MS run; tests as in **a–c**. Fractional abundances are corrected for natural isotope abundance, mass spectra normalised to internal standard and cell count. Data are mean ± s.e.m. Exact P values are shown.

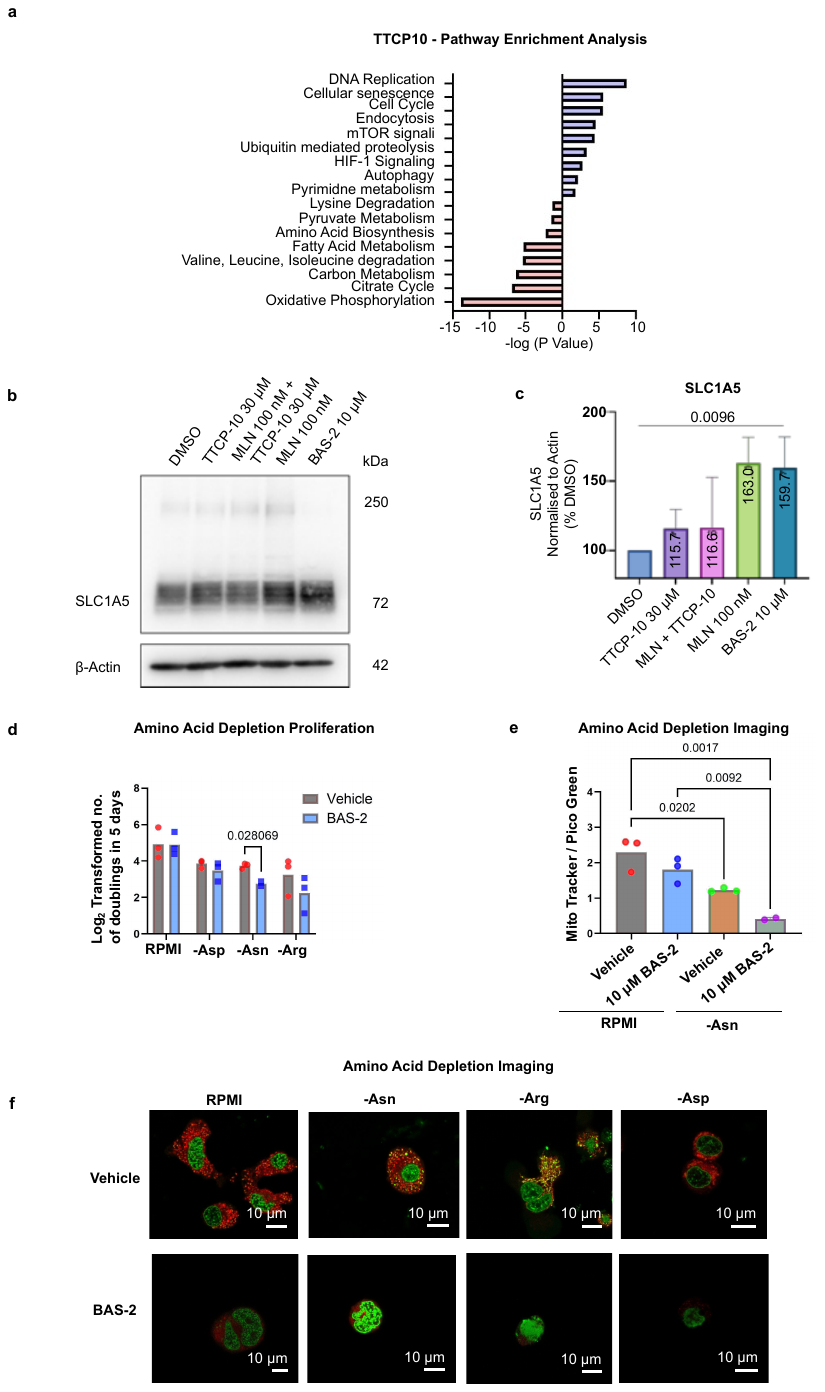

**Supplementary Fig. 3 | SLC1A5 regulation, amino-acid dependence and TTCP-10 pathway enrichment.**

**(a)** KEGG pathway enrichment of proteins changed by TTCP-10 in JJN3 cells. Positive values mark pathways enriched among upregulated proteins and negative values those among downregulated proteins (signed −logP) **(b)** Immunoblot of SLC1A5 in JJN3 cells treated for 24 h with DMSO, TTCP-10 (30 μM), MLN (100 nM) plus TTCP-10 (30 μM), MLN (100 nM) or BAS-2 (10 μM). Glycosylated (~72 kDa) SLC1A5 is marked; β-actin is the loading control. **(c)** Densitometry of SLC1A5 from a, normalised to actin and expressed as a percentage of DMSO; the mean of each condition is given within the bar and the exact P value is shown, n=3 biological replicates, one way Anova. **(d)** Population doublings over 5 days (log₂-transformed) of JJN3 cells in complete RPMI or in RPMI lacking aspartate (–Asp), asparagine (–Asn) or arginine (–Arg), with vehicle or 10 μM BAS-2. Bars show the mean, with individual replicates as points; n = 3 independent experiments; two-way ANOVA. **(e)** Mitochondrial content (MitoTracker relative to Pico Green) in JJN3 cells in RPMI or –Asn medium, with vehicle or 10 μM BAS-2. n = 3 independent experiments; one-way ANOVA with Tukey’s multiple comparisons test. **(f)** Representative confocal images for d across complete, –Asn, –Arg and –Asp media, with vehicle or BAS-2. MitoTracker is shown in red and Pico Green in green. Scale bars, 10 μm.

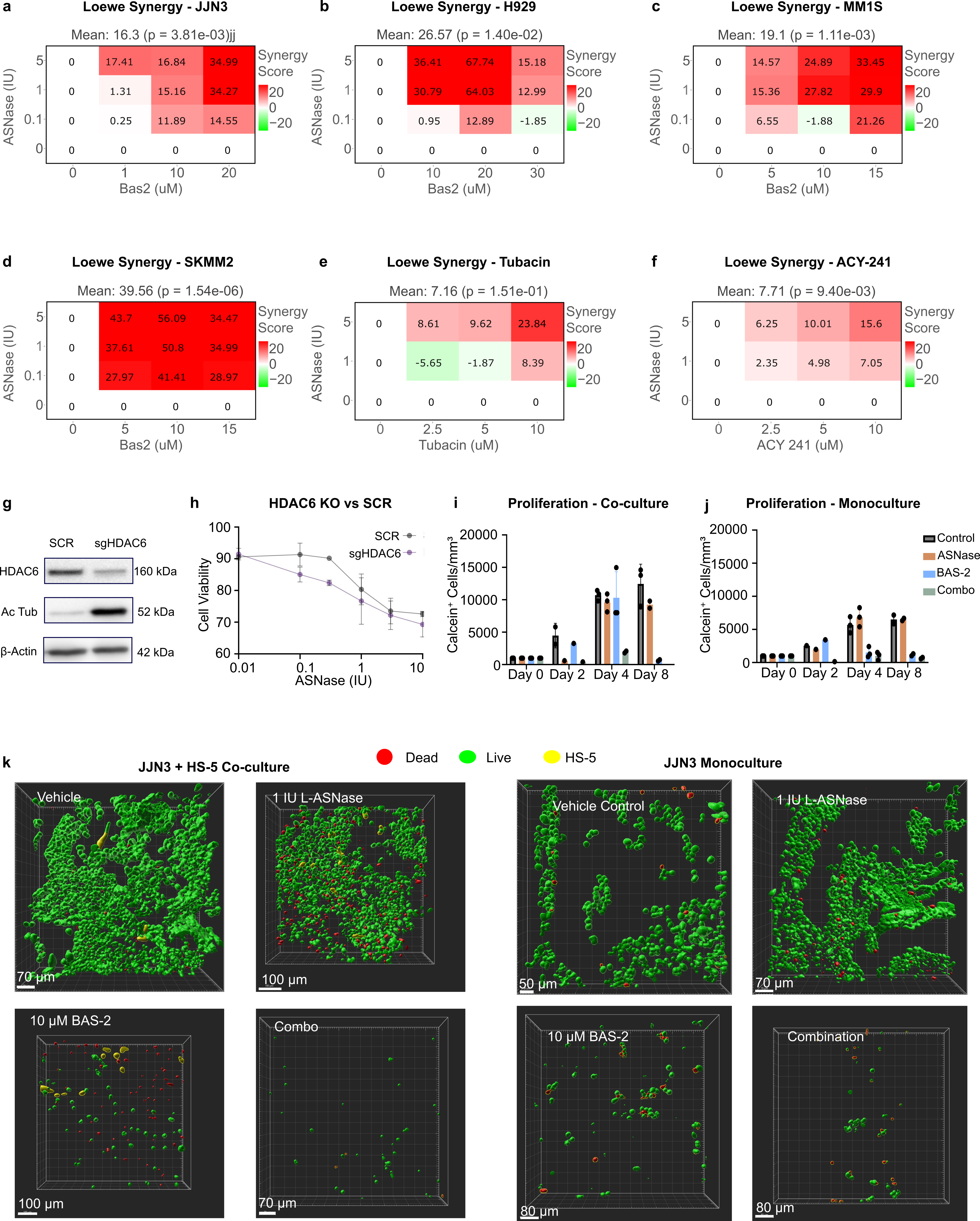

**Supplementary Fig. 4 | Genetic HDAC6 loss, 3D co-culture models and synergy across lines and inhibitors.**

**(a-f)** Loewe synergy maps for L-asparaginase combined with the HDAC6 inhibitors BAS-2 in JJN2 (**a**), H929 (**b**), MM1S (**c**), SKMM2 (**d**), L-Asn with tubacin (**e**) or ACY-241 (**f**) in JJN3 cells. In (**a-f**) values above 0 indicate synergy and values below 0 antagonism. Cells were seeded, treated with L-asparaginase 24 h later and with BAS-2 or HDAC6 inhibitor after a further 24 h, and viability was measured by Annexin V–PI staining after 24 h. The mean Loewe score and P value (SynergyFinder) are given above each map. n = 3 independent experiments. **(g)** Immunoblot of HDAC6 and acetylated α-tubulin (Ac tub) in JJN3 cells expressing scrambled (SCR) or HDAC6-targeting sgRNA (sgHDAC6). β-actin is the loading control. **(h)** Viability of SCR and sgHDAC6 JJN3 cells after L-asparaginase (0.01–10 IU, log scale) for 24 h. Data are mean ± s.e.m.; n = 3 biological replicates. **(I,j)** Calcein+ cell density (cells per mm³) in 3D bioprinted scaffolds of JJN3 cells co-cultured with HS-5 – mcherry tagged stromal cells (**i**) or JJN3 alone (**j**). Scaffolds were treated 24 h after seeding with vehicle, 1 IU L-asparaginase, 10 μM BAS-2 or the combination, and imaged on days 0, 2, 4 and 8 post treatment. n = 2-3 scaffolds across 2 experiments. **(k)** Representative 3D renderings for c and d at day 8. Live (Calcein^+^) cells are green, dead (PI⁺) cells red and HS-5 cells yellow. Scale bars are given in each panel.

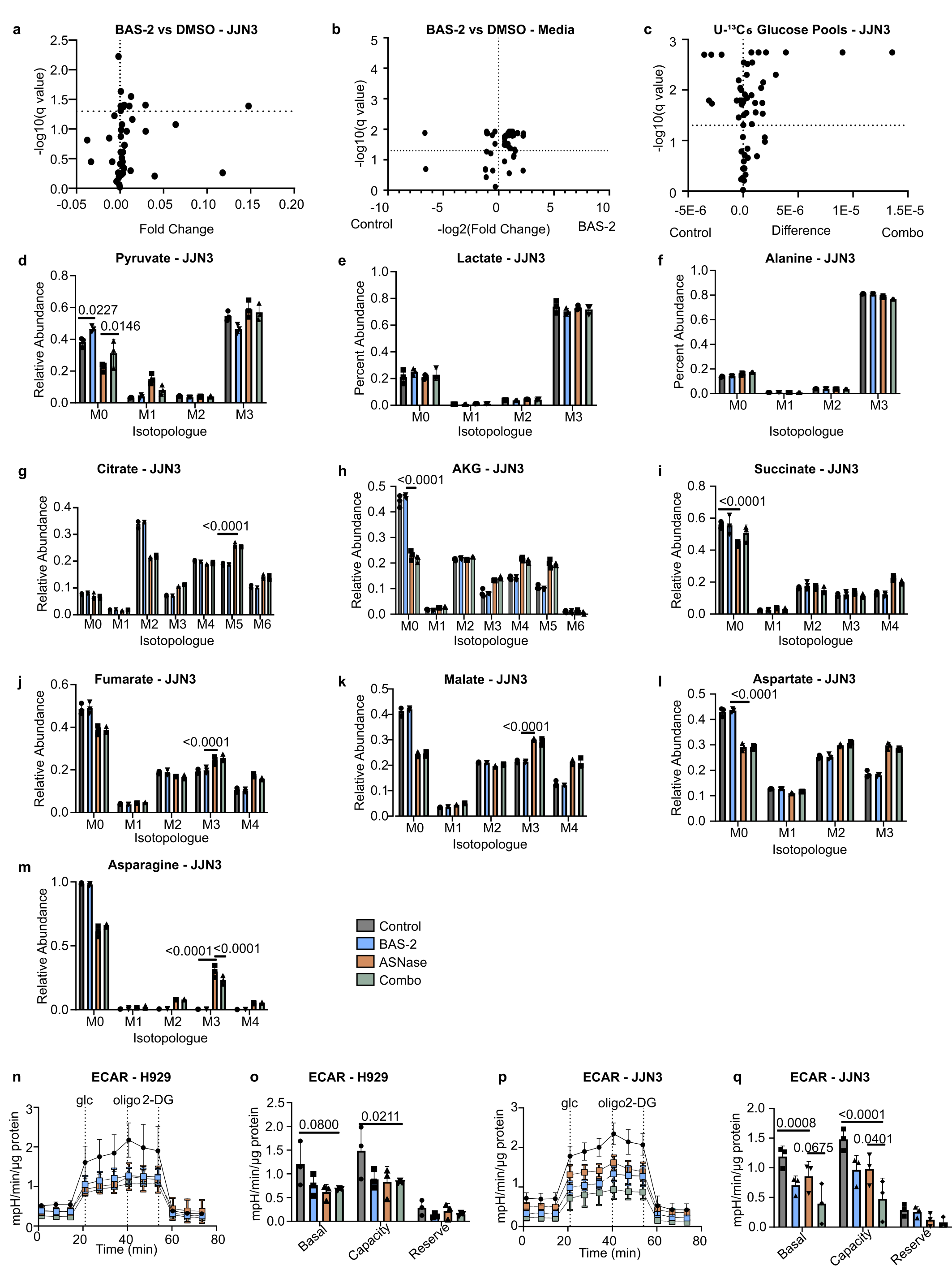

**Supplementary Fig. 5 | Metabolite pools, glucose-derived labelling and glycolytic flux under combination treatment.**

**(a)** Intracellular metabolite abundance in JJN3 cells treated with 10 μM BAS-2 versus DMSO. The dashed line marks q = 0.05. **(b)** Spent-media metabolites from the same treatment as a. **(c)** Total pool differences of U-¹³C_6_-glucose-derived metabolites in JJN3 cells, combination versus control. **(d–m)** U-¹³C_₆_-glucose isotopologue distributions of pyruvate (**d**), lactate (**e**), alanine (**f**), citrate (**g**), α-ketoglutarate (AKG; **h**), succinate (**i**), fumarate (**j**), malate (**k**), aspartate (**l**) and asparagine (**m**) in JJN3 cells treated with control, 10 μM BAS-2, 0.1 IU L-asparaginase or the combination. L-asparaginase was added 24 h and BAS-2 48 h after seeding, followed by 24 h in tracing medium. n = 3 independent cell cultures, analysed in a single GC–MS run; ordinary two-way ANOVA. **(n–q)** Extracellular acidification rate (ECAR) during a glycolysis stress test in H929 (**n,o**) and JJN3 (**p,q**) cells under the four treatment conditions. Traces (**n,p**) and derived basal glycolysis, glycolytic capacity and reserve (**o,q**) are shown, normalised to protein. glc, glucose; oligo, oligomycin; 2-DG, 2-deoxyglucose. n = 3 independent experiments; two-way ANOVA. Data are mean ± s.e.m.

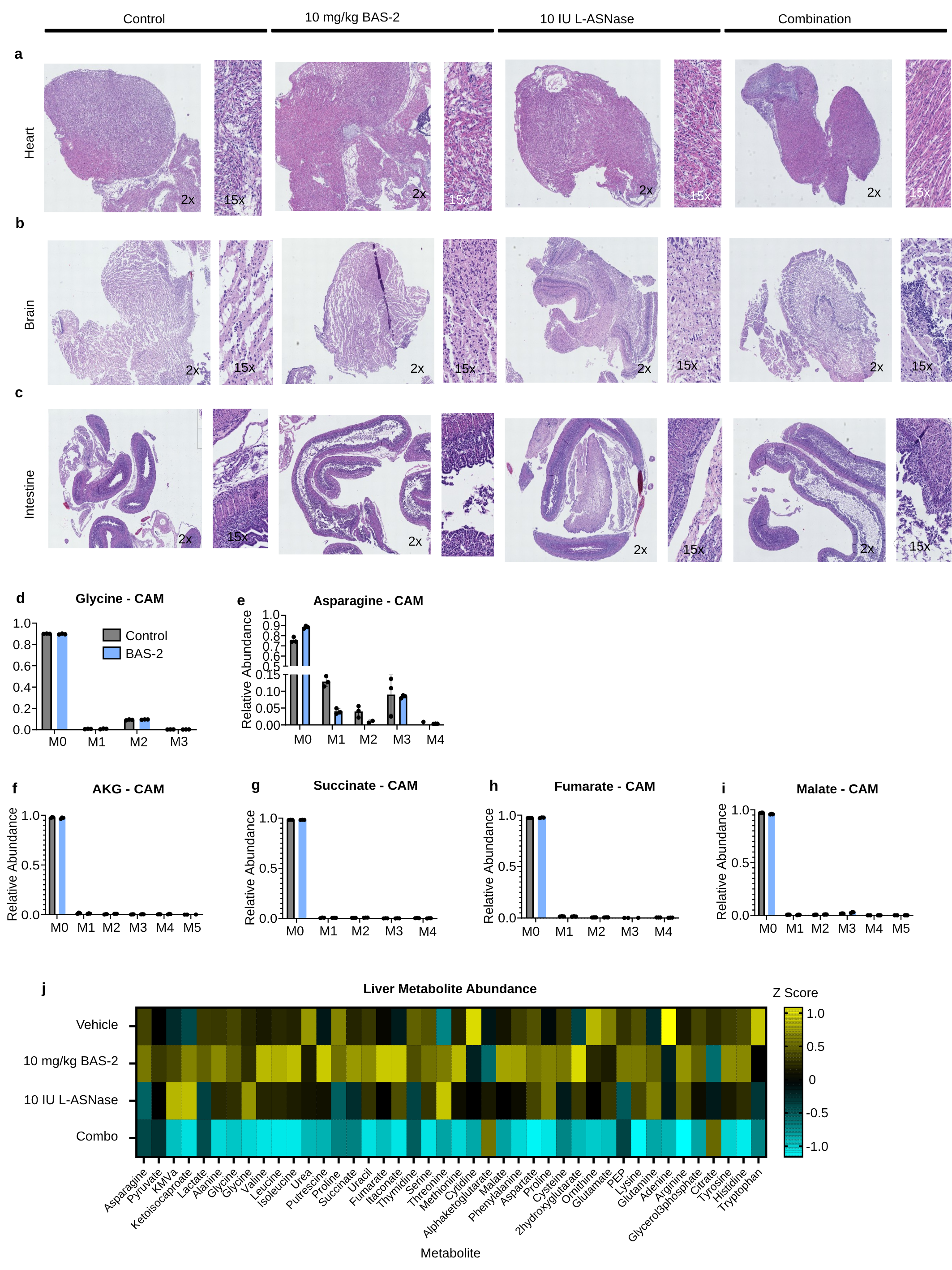

**Supplementary Fig. 6 | Organ histology, tumour tracing and liver metabolomics in the CAM model.**

**(a–c)** Representative H&E sections of chick embryo heart (**a**), brain (**b**) and intestine (**c**) after treatment with control, 10 mg/kg BAS-2, 10 IU L-asparaginase or the combination. Images are shown at 2× and 15× magnification. n = 3 embryos per condition. **(d–i)** U-¹³C_₆_-glucose isotopologue distributions of glycine (**d**), asparagine (**e**), α-ketoglutarate (AKG; **f**), succinate (**g**), fumarate (**h**) and malate (**i**) in JJN3 CAM tumours from control or BAS-2-treated embryos. n = 3 tumours, analysed in a single GC–MS run; one unpaired two-tailed t-test per isotopologue. Data are mean ± s.e.m. **(j)** Heat map of embryo liver metabolite abundance (row z-scores) across the four treatment conditions. n = 3 livers per condition.

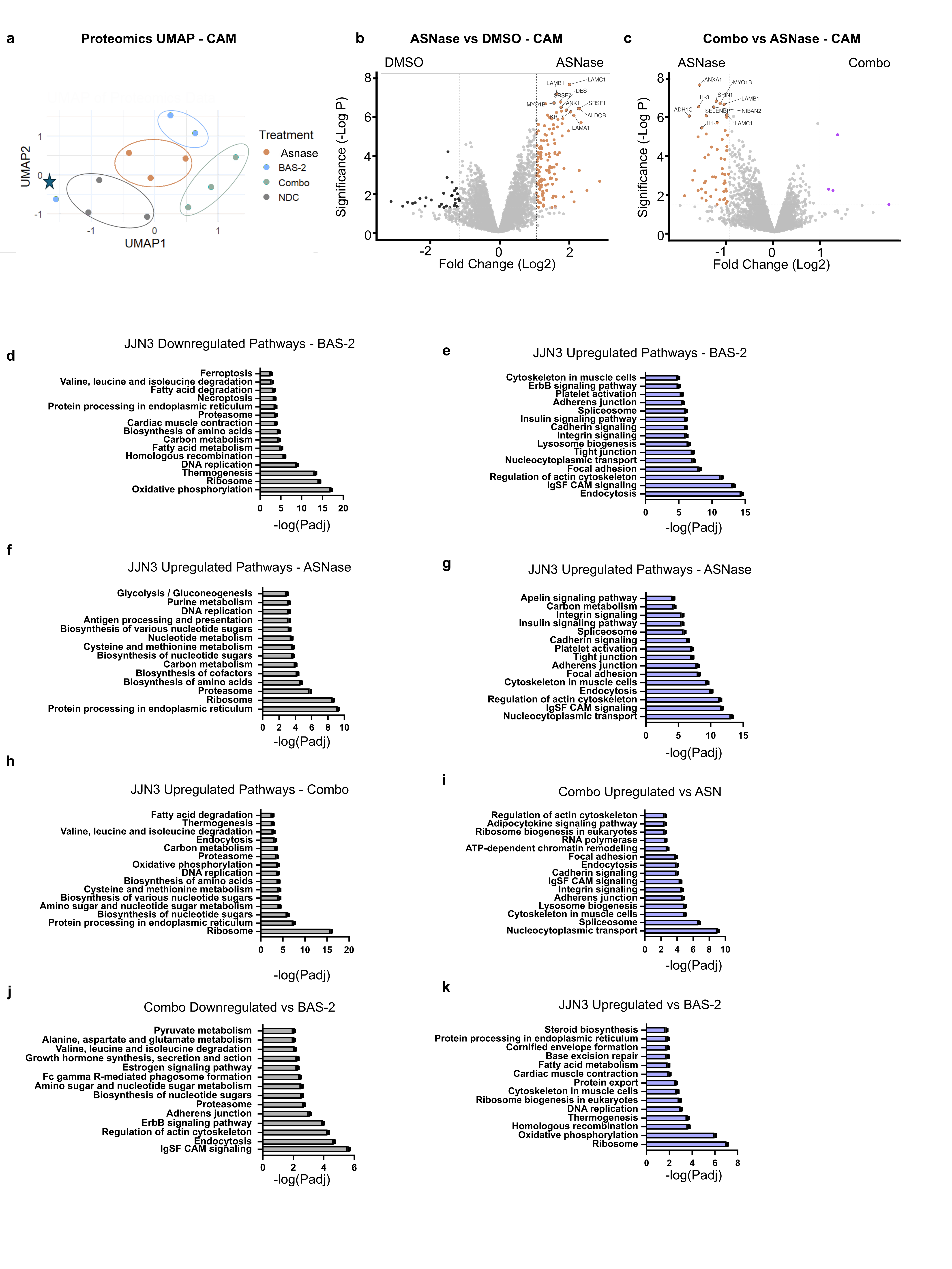

**Supplementary Fig. 7 | Proteomic response of CAM tumours and JJN3 cells to single agents and the combination.**

(**a**) UMAP of whole-tumour proteomes from JJN3 CAM xenografts treated with no-drug control (NDC), L-asparaginase (ASNase), BAS-2 or the combination. Ellipses group the replicates of each condition. The star marks an outlier sample. **(b,c)** Volcano plots of CAM tumour proteomes: L-asparaginase versus DMSO (b) and combination versus L-asparaginase (**c**). Dashed lines mark the fold-change and P thresholds; selected proteins are labelled. **(d–k)** KEGG pathways enriched among JJN3 proteins downregulated (grey) or upregulated (blue) after BAS-2 (**d,e**), L-asparaginase (**f,g**), combination versus L-asparaginase (**h,i**) and combination versus BAS-2 (**j,k**). Enrichment is shown as −log_₁₀_ adjusted P. n = 3 per condition.

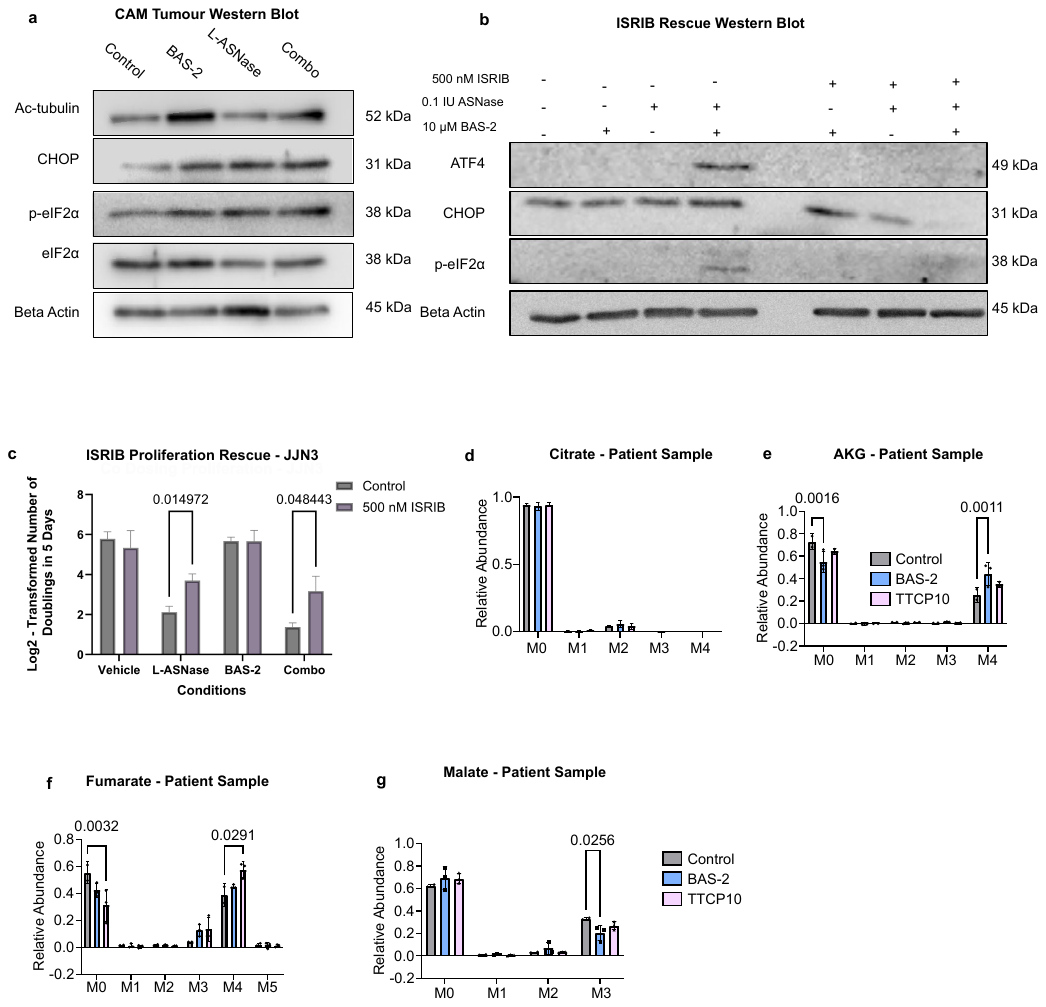

**Supplementary Fig. 8 | Integrated stress response activation, ISRIB rescue and primary-sample tracing.**

**(a)** Immunoblot of ATF4, CHOP and phosphorylated eIF2α (p-eIF2α) in JJN3 cells treated with 10 μM BAS-2, 0.1 IU L-asparaginase or both, with or without 500 nM ISRIB. Cells were seeded, treated 24 h later with L-asparaginase, then 24 h later with BAS-2 with or without ISRIB, and harvested after a further 24 h (72 h in total). β-actin is the loading control. **(b)** Immunoblot of acetylated α-tubulin, CHOP, p-eIF2α and total eIF2α in JJN3 CAM tumours from each treatment group. β-actin is the loading control. Blots in a and b are representative of n = 2 biological replicates. **(c)** Population doublings over 5 days (log_₂_-transformed) of JJN3 cells treated with vehicle, L-asparaginase, BAS-2 or the combination, with or without 500 nM ISRIB. Data are mean ± s.e.m.; n = 3 independent experiments; two-way ANOVA. **(d–g)** Fractional isotopologue distributions of citrate (**d**), α-ketoglutarate (AKG; (**e**), fumarate (**f**) and malate (**g**) in CD138⁺ primary myeloma cells labelled with U-¹³C₆-glucose and treated with vehicle, BAS-2 or TTCP-10 for 16 h. Cells come from one patient and were cultured in three separate plates (n = 3 technical replicates), analysed in a single GC–MS run. Ordinary two-way ANOVA. Data are mean ± s.e.m.; exact P values are shown.

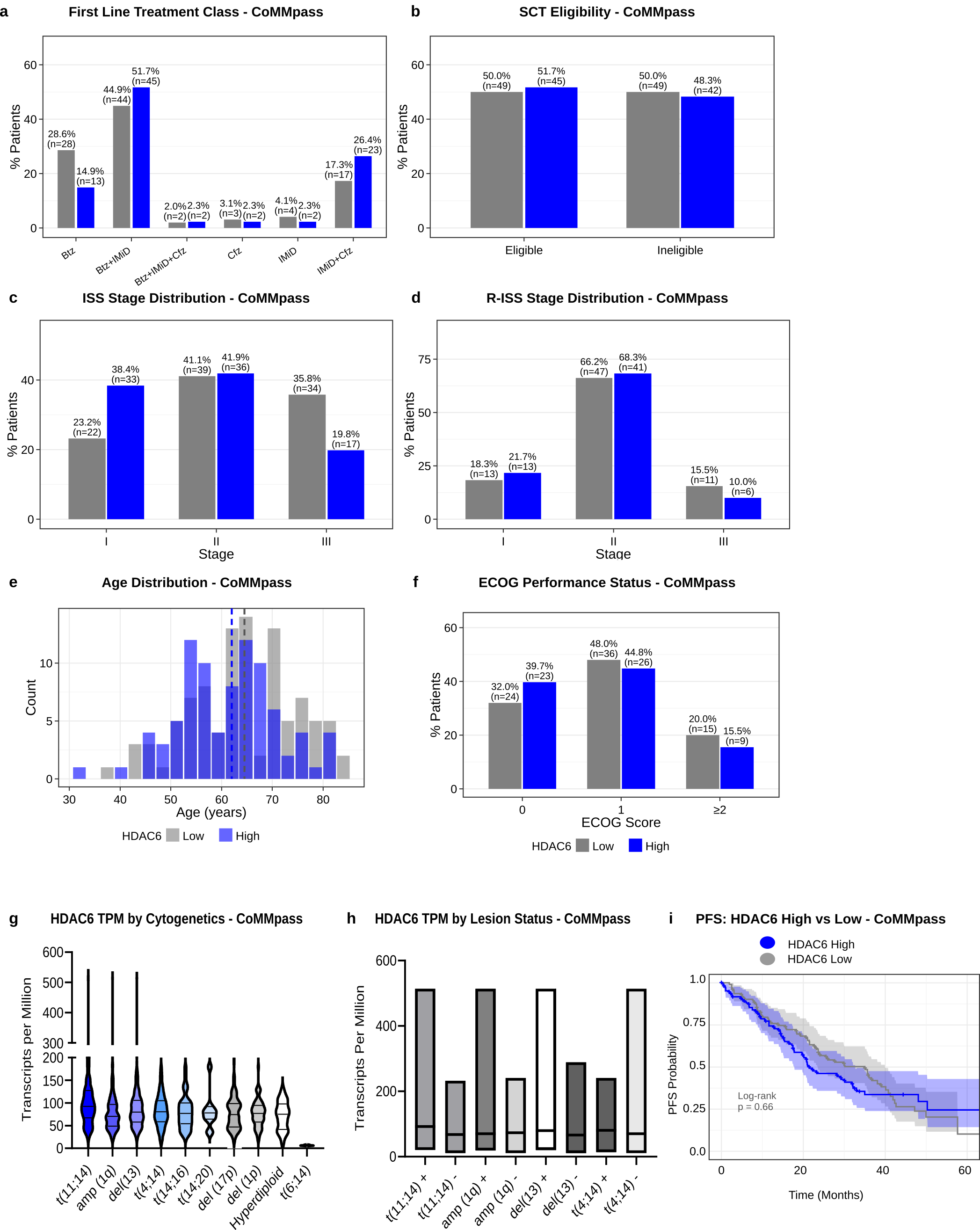

**Supplementary Fig. 9 | Clinical characteristics and HDAC6 expression in the CoMMpass HDAC6-high and HDAC6-low cohorts.**

HDAC6-high tumours are shown in blue and HDAC6-low tumours in grey (top and bottom 100 by HDAC6 TPM; MMRF CoMMpass IA14) **(a)** First-line treatment class. Btz, bortezomib; IMiD, immunomodulatory drug; Cfz, carfilzomib. **(b)** Stem-cell transplant (SCT) eligibility. **(c,d)** ISS (c) and R-ISS (d) stage at diagnosis. **(e)** Age at enrolment; dashed lines mark the group medians. **(f)** ECOG performance status. In a–d and f, bars show the percentage of patients with a known value; patient numbers are given above each bar. **(g)** HDAC6 expression (TPM) across all CoMMpass patients carrying each cytogenetic abnormality. Violins show the distribution, with the median and quartiles marked. The y axis is broken at 200 TPM. **(h)** HDAC6 TPM in patients positive (+) or negative (–) for t(11;14), amp(1q), del(13) and t(4;14). Floating bars span the minimum to maximum, with a line at the median. **(i)** Kaplan–Meier estimates of progression-free survival for HDAC6-high (n = 87) and HDAC6-low (n = 97) patients. Shading shows 95% confidence intervals, and the number at risk is given below. Patients were included if outcome data were available from the original selection. Log-rank P = 0.66.

**Supplementary Methods:**

**Lentiviral Transfections**

Lentiviral CRISPR constructs were pre-generated in-house: sgRNAs targeting *HDAC6* and a non-targeting scramble control (sgSCR) had been cloned into pLentiCRISPRv2 and sequence-verified, and the corresponding transformed Stbl3 clones archived as glycerol stocks at −80 °C.

For each construct, glycerol stocks were streaked or inoculated directly into 6 ml LB containing ampicillin (100 µg/ml) and cultured for 16 h at 37 °C, 210 rpm. Plasmid DNA was isolated by maxiprep (Qiagen), quantified by spectrophotometry, and molecular weight-validated by gel electrophoresis.

Lentivirus was produced in low-passage HEK293T cells (4 × 10⁶ per 10 cm dish) by co-transfection of 12 µg sgHDAC6 or sgSCR plasmid with 5 µg psPAX2 and 2.1 µg pMD2.G using TransIT-LT1 (Mirus) in Opti-MEM. Viral supernatants were harvested at 48 and 72 h, pooled, filtered through a 0.45 µm PVDF membrane and supplemented with 8 µg/ml polybrene. JJN3 cells (1 × 10⁶ per 10 cm dish) were transduced for 4 h, after which virus was replaced with fresh medium. Puromycin selection was applied for 6 days at a concentration predetermined by kill curve (0–10 µg/ml) for each line. Knockout was validated by immunoblot, and selected lines were maintained in low-dose puromycin (≤0.2 µg/ml).

**siRNA-mediated HDAC6 knockdown**

Cells were seeded in 6-well plates at approximately 40–50% confluency at the time of transfection. siRNA transfections were performed using Lipofectamine RNAiMAX according to the manufacturer’s instructions. siRNA was diluted in Opti-MEM and combined with Lipofectamine RNAiMAX diluted separately in Opti-MEM. Complexes were incubated for 10–20 min at room temperature prior to addition to cells. Cells were transfected with siRNA at a final concentration of 20 nM in a total volume of 2 mL per well, with 5 μL RNAiMAX in each well. Following transfection, cells were incubated for 48–72 h prior to downstream analyses. Non-targeting siRNA controls were included in all experiments. 20 nM siRNA and same v/v ratio of RNAiMAX was used for all experiments, regardless of well volume.

**CoMMpass analysis**

5.6.1	Data acquisition and pre-processing

Bulk RNA sequencing data and associated metadata were obtained from the Multiple Myeloma Research Foundation (MMRF) CoMMpass Study (release IA14a) ^1^. Gene-level normalised TPM matrices were imported into R (v4.4.1) and subjected to standard quality control procedures. Low-abundance transcripts were filtered out.

5.6.2	HDAC6 expression stratification

Normalised expression values for HDAC6 were used to stratify samples. To generate maximally contrasting groups, the 100 samples with the highest HDAC6 expression (HDAC6-high) and the 100 samples with the lowest expression (HDAC6-low) were selected for downstream analyses. Samples with intermediate expression values were excluded from group-based comparisons.

5.6.3	Differential gene expression analysis

Differentially expressed genes (DEGs) between the HDAC6-high and HDAC6-low groups were identified using limma, employing a linear modelling framework to compare between cohorts. DEGs were defined as genes with an adjusted P value (P_adj_) < 0.05 and an absolute log_2_ fold change > 1

5.6.4	Pathway enrichment analysis

Functional enrichment analysis was conducted using clusterProfiler. Gene Ontology (GO) Biological Process and KEGG pathway enrichment analyses were performed on significantly upregulated and downregulated DEGs separately ^2,3^. Over-representation tests were conducted using background gene sets matched to the filtered transcriptome. Pathways with FDR < 0.05 were considered significantly enriched.

5.6.5	Clinical metadata and survival outcome extraction

Clinical and demographic variables were extracted from the CoMMpass IA14 per-patient dataset (MMRF_CoMMpass_IA14_PER_PATIENT), including age at enrolment, sex, ISS stage, R-ISS stage, ECOG performance status, SCT eligibility, and first-line treatment class. Progression-free survival (PFS) were derived from the MMRF_CoMMpass_IA14_STAND_ALONE_SURVIVAL dataset. The pre-computed time-to-event columns within CoMMpass (ttos, ttpfs) are populated exclusively for patients who experienced the relevant event and are absent for censored observations; time variables were therefore constructed manually. PFS was defined as days from treatment start to first confirmed progression or death (censpfs = 1; event day: pfsdy), and to the last follow-up date (lastdy, or lvisitdy where lastdy was missing) for censored patients. All time variables were converted to months.

*Cytogenetic data extraction and recoding*

Cytogenetic abnormalities were extracted from the per-patient visit dataset (MMRF_CoMMpass_IA14_PER_PATIENT_VISIT). The D_TRI_cf flag was used to identify visits at which conventional cytogenetics or FISH had been performed; only the earliest such visit per patient was retained. Abnormality flags (D_TRI_abn1, abn4, abn6, abn8, abn10, abn12) employ a presence-only encoding: a value of 1 indicates the abnormality was detected, and a missing value in a tested patient indicates the abnormality was absent. Binary variables were constructed accordingly, with absent coded as FALSE and present as TRUE. Patients lacking any cytogenetic testing record were excluded from the denominator for each feature and retained as NA. The following abnormalities were assessed: del(13) (abn1), del(17p) (abn4), amp(1q) (abn6), t(4;14) (abn8), t(11;14) (abn10), and t(14;16) (abn12). Hyperdiploidy was classified as present when Hyperdiploid == “Yes” and absent when D_TRI_trisnone == 1 (no trisomies detected); all other patients were treated as untested.

*Survival and confounder analyses*

Survival analyses were performed in R (v4.4.1) using the survival and survminer packages. PFS was estimated by the Kaplan–Meier method and compared between HDAC6-high and HDAC6-low groups by log-rank test. Unadjusted and multivariable Cox proportional hazards models were fitted for both endpoints; covariates in the primary adjusted model were age, sex, ISS stage, SCT eligibility, and serum creatinine, selected on the basis of ≥98% non-missing data within the cohort. ECOG performance status was included in a pre-specified sensitivity model (approximately 72% covariate coverage). Results are expressed as hazard ratios (HR) with 95% confidence intervals (CI).

Baseline characteristics were summarised across HDAC6-high and HDAC6-low groups using the tableone package, with standardised mean differences (SMDs) reported to assess covariate balance (SMD > 0.1 considered indicative of potential imbalance). ISS and R-ISS stage distributions were compared by chi-squared test. Cytogenetic abnormalities were assessed by Fisher’s exact test; p-values were adjusted for multiple testing using the Benjamini–Hochberg procedure. Due to sparse cytogenetic data (3–53 events per feature across 184 tested patients), cytogenetic variables were not included in multivariable Cox models. Bar charts display the percentage of tested patients positive for each feature, with absolute counts labelled within bars. All tests were two-sided; P < 0.05 was considered significant.

**Flow cytometry**

*Annexin V–FITC / PI–PE viability assay*

Cell viability was evaluated using Annexin V–FITC and Propidium Iodide–PE staining. Following experimental treatment, cells were collected, washed in phosphate-buffered saline (PBS), and incubated with Annexin V–FITC (0.25 mg/ml, BioLegend) together with PI–PE (.5 mg/ml, Sigma) in annexin/PI buffer (10 mM HEPES, 140 mM NaCl, 2.5 mM CaCl_2_). Stained samples were analysed by flow cytometry on a Beckman Coulter CytoFLEX LX using standard settings for FITC and PE fluorescence and a BD FACS Canto II using standard settings. 10,000 events from the single-cell gated population were collected per sample. Data was processed in FlowJo, where debris and doublets were excluded prior to determining the proportion of viable cells (Annexin V^-^ / PI^-^).

*Protein synthesis assay (OPP incorporation)*

Protein synthesis was quantified using the Cayman Chemical Protein Synthesis Assay Kit (Cat. 601100). Cells were incubated with O-propargyl-puromycin (OPP) to label newly synthesised proteins, washed, and fixed according to the manufacturer’s instructions. 1 × 10^5^ JJN3 cells were seeded in 96-well plates 72 h prior to analysis and treated according to the indicated dosing schedule ^4^.

Cells were collected by centrifugation (500 × g, 5 min) and resuspended in RPMI medium containing OPP at a 1:250 dilution, followed by incubation for 30 min at 37°C. Cells were subsequently washed and fixed using fixation buffer for 5 min at room temperature. Following fixation, cells were washed with washing buffer and subjected to click chemistry labelling using a reaction mixture containing TBS, copper sulfate, 5-FAM azide, and ascorbic acid. Cells were incubated in the reaction mixture for 30 min at room temperature protected from light. Following labelling, cells were washed and resuspended in TBS prior to analysis by flow cytometry. OPP incorporation, indicative of nascent protein synthesis, was quantified using the FITC channel.

**Drug synergy quantification**

Synergy between BAS-2 and L-ASNase, and between alternative HDAC6 inhibitors (ACY-241, Tubacin) and L-ASNase, was assessed using full dose-response matrices. BAS-2 was titrated across a range of 0–30 μM (variable per cell line); ACY-241 and Tubacin were titrated across a range of 0–10 μM. L-ASNase was titrated at 0, 0.1, 1.0, and 5.0 IU/ml. Cells were dosed following the standard treatment scheme described above and viability assessed by Annexin V–FITC / PI–PE flow cytometry at Day 3. Viability matrices were exported and synergy scores computed in SynergyFinder using the Loewe additivity model^5^.

**Amino acid exclusion screen**

*Amino acid exclusion media preparation*

Amino acid-free RPMI powder (US Biological, 38210000) was reconstituted according to the manufacturer’s instructions and adjusted to pH 7.2 ± 0.1. Individual amino acids were weighed based on the standard RPMI formulation, and 19 of 20 amino acids were added back for each exclusion condition (L-amino acids kit, Supelco LAA21-1KT). Media were supplemented with 10% FBS and 1% penicillin–streptomycin, then sterile-filtered using a 0.2 μm syringe filter (Corning, CLS431229).

*Annexin V / PI viability arm*

JJN3 cells (50,000 per condition) were pelleted at 500 × g for 5 min, washed once in PBS, and resuspended in amino acid-depleted RPMI for 24 h. Cells were then treated with BAS-2 (10 or 20 μM) or 0.1% DMSO for an additional 24 h. Viability was assessed by Annexin V/propidium iodide staining followed by flow cytometry as described above.

*Proliferation arm*

For the focused proliferation arm in -Arg, -Asn, and -Asp conditions, JJN3 cells were prepared and resuspended in the relevant amino acid-depleted RPMI as described above, then treated with vehicle or 10 μM BAS-2 and followed across the duration of the 2D Proliferation Assay protocol below. 10,000 cells were seeded into 24 well plates.

**5.11	Spent medium amino acid measurement**

Conditioned media samples were collected prior to intracellular metabolite extraction and stored at -80°C until processing. Metabolite extraction was carried out as described by Cordes and Metallo. or metabolite extraction, samples were thawed on ice and mixed gently by inversion^6^. A 5 μL aliquot of media was transferred to a fresh Eppendorf tube containing 90 μL dH_2_O and 0.2 μg norvaline as an internal standard. Samples were maintained on ice throughout processing.

Extraction was performed by sequential addition of 250 μL cold methanol followed by 250 μL chloroform. Samples were incubated on a shaker at 4°C for 5 min and centrifuged at 10,000 × g for 5 min at 4°C to induce phase separation. The upper aqueous phase containing polar metabolites was transferred to a fresh tube and evaporated to dryness under nitrogen prior to downstream GC/MS analysis. Fresh media was harvested in tandem for comparison. Samples were frozen at -80°C until being run on GC-MS. Refer to GC-MS Tracing section for further details.

**Confocal and STED microscopy**

*MitoTracker Deep Red and Pico Green staining*

JJN3 cells were prepared for mitochondrial and DNA content imaging using concurrent staining with MitoTracker Deep Red FM and PicoGreen. Cells were washed in PBS and incubated with both dyes simultaneously at manufacturer-recommended concentrations (200 nM MitoTracker Deep Red, 0.1% v/v PicoGreen) under standard culture conditions and protected from light for 30 minutes. Following incubation, cells were washed again in PBS prior to downstream analysis. Fluorescence intensities from MitoTracker Deep Red and PicoGreen were used to evaluate relative mitochondrial content and nuclear DNA levels, respectively.

*PKMO and PicoGreen staining*

JJN3 cells were prepared for mitochondrial structure imaging using concurrent staining with PKMO and PicoGreen. Cells were washed in PBS and incubated with both dyes simultaneously, protected from light, and maintained under standard culture conditions for 30 minutes (100 nM PKMO, 0.1% v/v PicoGreen). After incubation, cells were washed again in PBS and prepared for STED microscopy. PKMO and PicoGreen fluorescence signals were used to assess mitochondrial structure and nuclear DNA content, respectively.

*Live-cell imaging slide preparation*

Ibidi μ-Slide 8-Well chambers were coated with poly-L-lysine (50 μg/mL in borate buffer) by adding 180–200 μL per well and incubating overnight at 37 °C. Slides were washed twice with distilled water and twice with PBS prior to cell seeding. JJN3 cells were counted, and 50,000 cells were seeded per well. After pelleting at 1000 rpm for 4 min, cells were incubated with staining solution in warm media for 30 min in the dark. Wells were washed three times with warm media and then resuspended in 500 μL warm media for imaging.

*Microscope setup*

A Leica Stellaris 8 STED 3D Falcon system equipped with a DMi8 inverted microscope and LASX software (version 4.6.0.2709) was used for all confocal and STED acquisitions. A variable white light laser (WLL, Leica) was used for excitation and a 775 nm STED laser for depletion. Three hybrid detectors (HyD S1, HyD S2, HyD X3; Leica) were used to collect signal, with HyD S2 used for all STED acquisitions. The microscope was equipped with an oil-immersion 100× objective (HC PL APO CS2, NA 1.4) and a 20× dry objective (NA 0.75). All confocal and STED images were acquired using the 100× objective unless otherwise stated. Brightfield images were simultaneously acquired using the Trans-PMT mode of a non-STED channel.

*Imaris Setup*

Z-stack images were imported into Imaris and converted to .ims files. Surfaces were created separately in the FITC (Pico Green) and PE (PKMO / Mitochondria) channels using the following parameters: surface detail 3.03 μm; diameter of largest sphere in object 11.4 μm; threshold 1.25 × 10^10^ (live channel) and 515 (dead channel). Each surface was mapped and geometric statistics extracted as described above.

**2D proliferation assay**

JJN3 cells (2.5 × 10^10^ per well) were seeded in 1 ml RPMI in 24-well plates in triplicate. After 24 h, cells were treated with BAS-2, L-ASNase, the combination, or amino acid-depleted media as described in the Amino acid exclusion screen, and incubated for an additional 96 h. Final cell numbers were determined on Day 5 by trypan blue exclusion and haemocytometer counting. Relative cell proliferation was expressed as the log_2_ fold change in cell density.

**3D bioprinted bone marrow constructs**

*Bioprinting of cell-laden constructs*

Three-dimensional cell constructs were fabricated using the RASTRUM bioprinter (Inventia Life Science). Prior to printing, the instrument completed Start-of-Day greenlighting to confirm operational nozzle status (minimum 2A/2B channels). Printing reagents were retrieved according to the system-generated Picklist and consisted of the RASTRUM 96-Well Plate Start Plus kit (Px02.29PH matrix; F177, F240, F3, F32). All reagents were thawed at room temperature for approximately 40 minutes and combined by pipetting to ensure homogeneity before loading into the designated cartridge reservoirs according to Cartridge Map A.

A sterile 96-well imaging plate (ThermoFisher 167008) was inserted into the printer, and an inert hydrogel base layer was printed into each well. During this step, cells were prepared for model printing. One million JJN3 cells were harvested and pelleted according to standard centrifugation conditions, then resuspended directly into the appropriate biofunctional Activator reagent (F177) to generate the Activator Cell Suspension. The suspension was transferred into the cartridge in positions specified by Cartridge Map B. Following inert base printing, the cell-laden Activator suspension and complementary Bioink components were printed into the wells to generate 3D constructs. Constructs were maintained in culture by gently exchanging medium along the well wall to avoid disturbing the hydrogel.

*Compound dosing*

Twenty-four hours after printing (Day 1), constructs were dosed by adding treatments directly to each well to a final working volume of 250 μL.

*Longitudinal Calcein-AM / PI imaging*

Constructs were imaged at Days 2, 4, and 8 post-printing. At each timepoint, stock Calcein-AM and Propidium Iodide (PI) were thawed 1 h prior to use, and staining media were prepared at 1 μg/mL Calcein-AM and 1 μg/mL PI in fresh culture medium. RASTRUM culture medium was removed and replaced with 250 μL staining solution per well. Constructs were incubated for 30 minutes at 37 °C, after which the stain was removed, constructs were washed with 250 μL PBS, and resuspended in RPMI for imaging.

*Fluorescence imaging*

Imaging was performed on a Leica fluorescence confocal microscope equipped with environmental control (37 °C, 5% CO_2_). Plates were imaged using a 20× dry objective. Calcein-AM and PI fluorescence were acquired using FITC (Detector 2) and PE (Detector 3) channels in analogue mode. Scanning parameters included a speed of 600, image resolution of 256 × 256 pixels, and a Z-step of 0.75 μm (or Smart Z). Fast Live mode was used to optimise gain and illumination while preventing saturation. Z-stack boundaries were defined manually for each construct.

*Surface reconstruction modelling*

Surface modelling was performed in Imaris software. Z-stack images were imported into Imaris and converted to .ims files. Sample surfaces were created in the FITC and PE channels, with FITC corresponding to Calcein^+^ live cells and PE channel to PI^+^ dead cells. Each surface was mapped and geometric statistics were extracted.

*Proliferation and viability extrapolation*

Overall surface counts were taken from live and dead surfaces in each image. Calcein^+^ cells were normalised to z-stack volume and reported as cells per mm^3^ to capture organoid proliferation over the imaging window. Each condition and timepoint was treated as a technical replicate. Viability was measured as the ratio of Calcein^+^ to PI^+^ surfaces in a given region of interest.

**CAM xenograft modelling**

*Egg preparation and incubation*

CAM model was carried out as described by Cymer et al. ^7^. Fertilised chicken eggs were cleaned thoroughly with 70% ethanol and water, labelled, and placed horizontally in a humidified incubator at 37–38 °C. The position of the air sac was marked prior to incubation. CAM experimental days are reported relative to tumour engraftment, with Day 0 corresponding to embryonic development day 9 (EDD9).

*Albumin removal and windowing (EDD3–4)*

Under sterile conditions, albumin was withdrawn from the air sac end of each egg using a syringe fitted with a 21–23 G needle. The needle was inserted 1–2 mm into the shell, rotated 90°, and advanced parallel to the shell surface to withdraw 3–4 ml albumin depending on egg size. Eggs showing white albumin were discarded due to suspected contamination. The puncture site was sealed with surgical tape. A small shell window was then cut above the embryo, ensuring the membrane remained intact, and resealed with clear surgical tape before returning eggs to the incubator.

*Tumour xenograft implantation (Day 0 / EDD9)*

Tumour cells were prepared by pelleting and resuspending 1 × 10^6^ JJN3 cells in a Matrigel/RPMI mixture (1:1 v/v). Plastic incubation rings were sterilised in 100% ethanol and dried under a heat lamp. Eggs were briefly warmed, after which the shell window was enlarged to expose a highly vascularised region of the CAM. A sterile ring was placed onto the CAM surface, and 50 μL of the tumour–Matrigel suspension was carefully dispensed within the ring while avoiding air bubble introduction. Control eggs received Matrigel alone, vehicle, or other designated control suspensions. Windows were resealed, and eggs were returned to the incubator.

*Cohort design, randomisation, and exclusion criteria*

Each experimental group comprised 6–10 eggs. Eggs were randomised between treatment conditions following successful tumour engraftment. Exclusion criteria included embryonic death, failed engraftment, or contaminated albumin (white) at the windowing step. Surviving eggs were tracked throughout the dosing window, and harvest was blinded to treatment condition where possible.

*Drug administration (Day 3 EDD12 and Day 5 EDD14)*

Drug solutions were prepared in FBS-free RPMI, maintaining final DMSO concentrations below 1%. Windows were reopened, and 100 μL of drug solution was applied directly onto the tumour xenograft on Day 3 (Dosing 1 EDD12) and Day 5 (Dosing 2 EDD14). Doses were 115 μg (10mg/kg) BAS-2 and 10 IU L-ASNase, or the combination per egg.

*Tissue harvest and sample processing (Day 7 EDD 16)*

Eggs were placed on cotton pads, and the windows were opened fully to allow removal of the embryo. Tumour tissue and surrounding CAM were excised; tumour volume was measured by calipers prior to fixation. Tumours were then fixed in 10% neutral buffered formalin for at least 24 h, or snap frozen in liquid nitrogen. Embryos were dissected to collect major organs (brain, heart, lungs, liver, gizzard, spleen, kidneys) using sterile scissors and forceps. Organs were either fixed or snap-frozen and stored in cryovials for downstream analyses.

**Stable isotope tracing**

*Tracer treatments*

Isotope tracing was carried out as described by Cordes and Metallo.

JJN3 multiple myeloma cells were seeded at 125,000 cells per well in 6-well plates and allowed to equilibrate for 24 h before treatment. Cells were treated following the standard drug treatment scheme described above and then transferred into medium containing either U-^13^C_6_-glucose or U-^13^C_5_-glutamine. For pulse tracing of glucose- or glutamine-derived carbon, cells were exposed to the labelled tracer for 30 minutes. For steady-state pool measurements, cells were exposed to the labelled tracer for 24 hours.

*Harvesting and metabolite extraction*

Extraction and measurement were done as specified by Cordes and Metallo**.** At the end of tracer incubation, cells were transferred to pre-chilled Eppendorf Safe-Lock tubes, washed twice with ice-cold 0.9% saline, and pelleted at 500 × g for 5 min at 4 °C. Pellets were resuspended in 500 μL -80 °C methanol (containing internal standards where required), incubated on dry ice for approximately 1 min, and mixed with 200 μL Milli-Q water containing 0.2 μg norvaline. After brief vortexing, 500 μL chloroform was added, and samples were vortexed for 10 min at 4 °C. Samples were centrifuged at >12,000 × g for 10 min at 4 °C to achieve biphasic separation. The upper methanol–water phase (polar metabolites) and the lower chloroform phase (lipids) were collected into new tubes without disturbing the protein interphase.

*Drying and storage*

Extracts were evaporated to dryness under nitrogen or in a refrigerated speed vacuum. Complete dryness was confirmed prior to derivatisation. Dried samples were stored at -80 °C until GC–MS analysis. For each tracing experiment, n = 3 refers to three separate cell cultures per condition; all samples from a given experiment were derivatised and analysed together in a single GC–MS run.

**Derivatisation of polar metabolites**

Dried aqueous-phase metabolite extracts and dried spent-medium extracts were derivatised in two steps following Cordes & Metallo. Samples were first incubated with 20 μL of 2% (w/v) methoxyamine hydrochloride (MOX; Supelco 33045-U) in pyridine at 40 °C for 60 min with shaking to protect keto groups. Samples were then silylated with 15 μL of MTBSTFA containing 1% *tert*-butyldimethylchlorosilane (Regis Technologies 1-270143-200) at 40 °C for 30 min. Any precipitate was removed by brief centrifugation prior to GC–MS injection. MTBSTFA derivatives were selected for their stability and suitability for targeted analysis of organic acids, amino acids, and TCA cycle intermediates.

**GC instrument, column and run program**

Derivatised samples were analysed on an Agilent 7890A gas chromatograph fitted with an Agilent DB-35MS UI capillary column (30 m × 0.25 mm × 0.25 μm; J&W Scientific 122-3832 UI) coupled to an Agilent 5977B mass spectrometer. One microlitre was injected in splitless mode at 270 °C with helium as carrier gas at 1 mL/min. The oven was held at 100 °C for 1 min, ramped to 255 °C at 3.5 °C/min, then to 320 °C at 15 °C/min, and held at 320 °C for 3 min (total run time 54.6 min). The transfer line was held at 290 °C.

**Mass spectrometer parameters**

The Agilent 5977B mass spectrometer operated in electron impact (EI) ionisation mode at 70 eV, with the source held at 230 °C and the quadrupole at 150 °C. Spectra were acquired in scan mode covering *m/z* 50–650 to capture the targeted metabolite fragment ions used in this study (pyruvate, lactate, alanine, citrate, α-ketoglutarate, succinate, fumarate, malate, aspartate, glutamate, glutamine, asparagine, and the amino-acid panel). Fragment ions and retention times were taken from the targeted ion list of Cordes & Metallo. Norvaline (Sigma-Aldrich N7502) was added to the methanol–water extraction phase at 5 μg/mL as internal standard.

**AMDIS–MATLAB integration and isotopologue analysis**

Raw GC–MS files were inspected in AMDIS (<http://www.amdis.net>) to identify metabolite peaks and update expected retention times in the MATLAB library scripts. Integration was then performed in MATLAB R2025b using in-house scripts invoking integrate_carbon(‘<filepath>’, ‘<library script>’, 1), where the final argument enables natural-abundance correction. Integrated isotopologue distributions were exported as .csv and imported into Excel and GraphPad Prism. Fractional contribution of ^13^C-glucose-derived carbon to de novo biosynthesis was calculated as Mn/M0 isotopologue ratios within each condition (M+4 for glutamine; M+3 for aspartate and asparagine).

**Extracellular flux analysis (Seahorse)**

Glycolytic flux was assessed in JJN3 and H929 cells using the Seahorse XF Glycolysis Stress Test Kit (Agilent, 103020-100) on a Seahorse XFe96 analyser. Sensor cartridges were hydrated overnight in XF Calibrant solution at 37°C in a non-CO₂ incubator. Assay medium was prepared from Seahorse XF Base Medium (Agilent, 103334-100) supplemented with 1 mM L-glutamine and pH-adjusted to 7.4 with NaOH immediately before use.

Cell-Tak adhesive (Corning, VWR #734-1081) was prepared in 0.1 M sodium bicarbonate (pH 8.0) with 1 M NaOH and applied at 25 µL per well to XFe96 cell culture microplates. Plates were incubated at room temperature for at least 20 minutes, after which wells were washed twice with 200 µL ddH₂O.

Cells were harvested, counted, and pelleted by centrifugation. The pellet was washed in pre-warmed assay medium without resuspension, then resuspended to a density of 1×10⁵ cells per well in 180 µL assay medium. Cells were seeded into coated wells and centrifuged at 1,200 rpm for 3 minutes with the brake disengaged to promote uniform adhesion. Wells were inspected for monolayer uniformity and plates were transferred to the non-CO₂ incubator while compounds were loaded. Blank wells (four corner wells) received 180 µL assay medium without cells.

Compounds were prepared from kit stocks and loaded into injection ports at the following final concentrations: glucose (10 mM, Port A), oligomycin (1 µM, Port B), and 2-deoxyglucose (2-DG; 50 mM, Port C). The extracellular acidification rate (ECAR) was recorded across three basal measurement cycles prior to sequential injection of each compound. Assay was preformed immediately after finishing compound preparation. Following extracellular acidification experiment, media was removed from plate, with care taken not to detach cells. Protein concentration was quantified via BCA assay directly following seahorse assay.

**ISRIB rescue experiments**

To test whether the integrated stress response is causally required for combination-induced cell death, ISRIB was added to the cell culture plate at a final concentration of 500 nM concurrent with BAS-2 administration on Day 2 of the standard combination dosing scheme. ASNase (or vehicle) was added on Day 1 and cells were harvested on Day 3 for downstream readouts (Annexin V/PI flow cytometry for the ISRIB rescue viability arm; whole-cell lysis and Western blotting for the ISRIB rescue biochemical arm). ISRIB was reconstituted in DMSO and prepared in single-use aliquots stored at -20 °C; final DMSO concentration was maintained below 0.1% in all wells, including matched vehicle controls.

**In vivo ^13^C_6_-glucose tracing in CAM tumours**

To validate the glycolytic block imposed by BAS-2 in the tumour setting, U-^13^C_6_-glucose was administered topically to CAM-engrafted JJN3 tumours in the vehicle control and BAS-2 arms (n = 3 eggs per group) 24 h prior to harvest. A solution of 10 mg U-^13^C_6_-glucose in 100 μL serum-free RPMI was prepared freshly and applied directly onto the tumour xenograft through the shell window on Day 6 (EDD 15). Windows were resealed and eggs returned to the incubator following dosing.

At harvest (Day 7 / EDD 16), tumours were resected, snap-frozen in liquid nitrogen, and stored at -80 °C until metabolite extraction. Frozen tumours were thawed on ice and manually homogenised using a mortar and pestle. Homogenates were resuspended in ice-cold 0.9% NaCl and further disrupted by probe sonication on ice (5 × 10 s pulses) at the same instrument settings described under *CAM tumour proteomics*. Metabolites were then extracted by the methanol–water–chloroform biphasic method described under *Stable isotope tracing*, with norvaline added at 5 μg/mL as internal standard. Drying, derivatisation, and GC–MS analysis were performed as described under *GC–MS metabolomics*.

**Proteomics**

*Mass spectrometry proteome analysis of JJN3 cells*

JJN3 multiple myeloma cells were cultured in triplicate in 6-well plates at a seeding density of 5 × 10^5^ cells/well. Cells were treated with vehicle or 10 μM BAS-2 for 16 h, then lysed in RIPA buffer supplemented with complete protease inhibitor tablets (Sigma-Aldrich). Lysates were centrifuged at approximately 30,000 × g for 10 min at 4 °C, and supernatants were collected for downstream processing.

Proteins were prepared using a standard SP3 workflow. A total of 50 μg protein per sample was diluted in lysis buffer containing 4 M urea, 50 mM ammonium bicarbonate, and 5 mM calcium chloride. Samples were reduced with 0.2 M DTT for 15 min at room temperature and alkylated with 4 mM IAA for 15 min at room temperature in the dark. Hydrophilic and hydrophobic SP3 beads (5 μL each) were washed in MS-grade water and added to lysates on a KingFisher Duo Prime system along with Pierce Trypsin Protease, MS Grade (Thermo Fisher, 90058) at 0.5 μg/μL. Ethanol (80%) was used in adjacent wells for bead-washing steps. Samples underwent overnight processing (8 h digestion at room temperature followed by storage at 4 °C).

Peptides were eluted in 0.1% formic acid, dried by speed vacuum, and stored at -20 °C.

Mass Spectrometry (Bruker timsTOF Pro)

Prior to creating a data-independent acquisition (DIA) method, a pool of the combined trypsin digested samples was analysed using dda-PASEF. Tryptic peptides were resuspended in 0.1% trifluoroacetic acid, and each sample was loaded onto an Evosep tip. The Evosep One was configured to elute and separate the peptides using a standard chromatography method (30 samples per day)^8^. The mass spectrometer was operated in positive ion mode with a capillary voltage of 1350 V, a dry gas flow of 3 L/min and a dry temperature of 180 °C. All data was acquired in data-dependent acquisition mode with parallel accumulation serial fragmentation (dda-PASEF), in which trapped ions were selected for MS/MS. A scan range of 100–1700 m/z was used, with four PASEF MS/MS frames per MS scan and a cycle time of 0.53 s^9^.

The resulting dda-PASEF file was searched in FragPipe version 24 ^10,11^ in basic-search mode to create a spectral library. This library was used to generate a custom DIA window scheme with py_diAID, which was embedded into the timsTOF dia-PASEF workflow ^12^. All data was acquired using dia-PASEF. The dia-PASEF settings were variable mass window width, isolation window overlap 0, 40 MS/MS windows, 20 MS/MS ramps, cycle time 2.23 s, and mass range 302.5–1198.1 m/z ^13^.

Data analysis

Proteome samples acquired using dia-PASEF were analysed using DIA-NN 2.3.0 Academia (data-independent acquisition by neural networks) ^14^. A predicted spectral library was generated within DIA-NN (library-free mode) from the Homo sapiens UniProtKB Swiss-Prot (reviewed) database (downloaded 19 February 2026). Search settings for the total proteome included: match between runs on, cysteine carbamidomethylation as a fixed modification, N-terminal methionine excision enabled, N-terminal acetylation and methionine oxidation as variable modifications, a maximum of one missed cleavage, precursor charge states 2–4, neural network classifier in cross-validated mode, retention time-dependent cross-run normalisation, and library generation set to RT and IM profiling. All output was filtered at 1% FDR.

*Bioinformatic analysis – In Vitro*

Output label-free quantification (LFQ) values were imported into Perseus (v2.0.7.0) and transformed to log_2_. Normality of sample distributions was confirmed before filtering for proteins with at least three valid values in one group. Volcano plots were generated using an P Value of 0.05 using VolcaNoseR software ([https://huygens.science.uva.nl/VolcaNoseR](https://eur02.safelinks.protection.outlook.com/?url=https%3A%2F%2Fhuygens.science.uva.nl%2FVolcaNoseR&data=05%7C02%7Cmatthewhiller24%40rcsi.ie%7Cc67f7551189c4ac5dcfe08df464def93%7C607041e7a8124670bd3030f9db210f06%7C0%7C0%7C639306972698788861%7CUnknown%7CTWFpbGZsb3d8eyJFbXB0eU1hcGkiOnRydWUsIlYiOiIwLjAuMDAwMCIsIlAiOiJXaW4zMiIsIkFOIjoiTWFpbCIsIldUIjoyfQ%3D%3D%7C0%7C%7C%7C&sdata=JEdwlhSTBMSuWg5q7JwVAIckhIb1GdzdAMbNZL0CMKk%3D&reserved=0)). Enrichment was performed with Enrichr software and was based on Go Biological Process 2023 Human library and pathway ranked by the p-value of Fisher’s exact test. Differentially abundant proteins from BAS-2 and TTCP-10 treatments, versus DMSO control, were used to construct a protein-protein interaction network using STRING in Cytoscape. The two networks were then merged using Cytoscape to identify the differentially abundant proteins common to both treatment groups. Go Biological process enrichment was then overlaid on the merged network.

*CAM tumour proteomics*

CAM-engrafted JJN3 tumours from each treatment arm were resected at harvest (Day 7/ EDD 16), placed into pre-labelled cryotubes, and snap-frozen in liquid nitrogen. Frozen tumours were thawed on ice, washed once in ice-cold PBS, and resuspended in RIPA lysis buffer supplemented with complete protease and phosphatase inhibitors (Sigma-Aldrich). Tissues were manually homogenised with a pestle and further disrupted by probe sonication on ice (5 × 10 s pulses). Lysates were then processed as described under *Western blotting* (centrifugation, clarification), and protein concentration was quantified by BCA assay (Thermo Fisher). Each lysate corresponded to a single tumour, with no pooling across animals. Clarified lysates were stored at -80 °C until proteomic sample preparation, which was performed using the SP3 and dia-PASEF workflow described under *Proteomics (in vitro)*.

*Bioinformatic analysis – CAM*

Protein groups intensities were log_2_-transformed and filtered to retain proteins quantified in at least 9 of 12 samples. Remaining missing values were imputed via left downshift method. Column mean minus 1.8 multiplied by column standard deviations. Random value was then derived from a normal distribution with a width of 0.3 standard deviations. Differential abundance between groups was assessed in limma (v.3.62.2)R(v.4.4.2) using a means model over treatment group. Volcano plots were generated via ggplot2(v.4.0.1) R package using a P.Value of 0.05. Pathway enrichment was preformed via cluterProfiler (v.4.14.6).

**Immunohistochemistry and haematoxylin and eosin staining**

Harvested CAM tumours were fixed in 10% neutral buffered formalin for a minimum of 24 h, processed, and embedded in paraffin. Sections of 4 μm thickness were cut using a Leica RM2135 rotary microtome. Haematoxylin and eosin (H&E) staining was performed on the Dako CoverStainer (Agilent) according to the manufacturer’s standard programme and used for morphological assessment only.

For immunohistochemical detection of CD138, staining was performed on a Bond-III automated immunostainer (Leica Biosystems). Antigen retrieval was carried out on-board using Bond Epitope Retrieval Solution 1 (Leica, AR9961) for 20 min. The primary antibody anti-CD138 (clone IHC138; Cell Signaling Technology, #30501) was diluted 1:300 in Bond Primary Antibody Diluent (Leica, AR9352). Antibody binding was detected using the Bond Polymer Refine Detection Kit (Leica, DS9800) with DAB signal enhancement using Bond DAB Enhancer (Leica, AR9432). Sections were counterstained with haematoxylin and coverslipped. CD138-positive area was quantified in ImageJ using a consistent threshold applied across all sections and expressed relative to total tissue area per image. Three biological replicates per treatment condition were analysed.
